# Targeting A-Kinase Anchoring Protein 12 Phosphorylation in Hepatic Stellate Cells Regulates Liver Injury and Fibrosis in Mouse Models

**DOI:** 10.1101/2022.03.15.484391

**Authors:** Komal Ramani, Nirmala Mavila, Aushinie Abeynayake, Maria Lauda Tomasi, Jiaohong Wang, Mitchitaka Matsuda, Ekihiro Seki

## Abstract

Trans-differentiation of hepatic stellate cells (HSCs) to activated state potentiates liver fibrosis through release of extracellular matrix (ECM) components, distorting the liver architecture. Since limited antifibrotics are available, pharmacological intervention targeting activated HSCs may be considered for therapy. A-kinase anchoring protein 12 (AKAP12) is a scaffolding protein that directs protein kinases A/C (PKA/PKC) and cyclins to specific locations spatiotemporally controlling their biological effects. It has been shown that AKAP12’s scaffolding functions are altered by phosphorylation. In previously published work, observed an association between AKAP12 phosphorylation and HSC activation. In this work we demonstrate that AKAP12’s scaffolding activity towards the endoplasmic reticulum (ER)-resident collagen chaperone, heat-shock protein 47 (HSP47) is strongly inhibited by AKAP12’s site-specific phosphorylation in activated HSCs. CRISPR-directed gene editing of AKAP12’s phospho- sites restores its scaffolding towards HSP47, inhibiting HSP47’s collagen maturation functions and HSC activation. AKAP12 phospho-editing dramatically inhibits fibrosis, ER stress response, HSC inflammatory signaling and liver injury in mice. Our overall findings suggest a pro-fibrogenic role of AKAP12 phosphorylation that may be targeted for therapeutic intervention in liver fibrosis.

## Introduction

Hepatic stellate cells (HSCs) constitute approximately 5-8% of the normal liver and are major sites for vitamin A storage in the body (1). During chronic liver injury, HSCs acquire a pro-fibrogenic phenotype or activated state that is critical in the liver’s response to injury (2). HSC activation causes increased production of extracellular matrix (ECM) components such as collagens and α-smooth muscle actin (α-SMA). Persistent injury leads to fibrosis due to abnormal accumulation of ECM (3). HSC pathways that cause fibrogenic responses in the liver can be targeted for therapeutic intervention.

Collagen maturation and secretion is facilitated by the endoplasmic reticulum (ER)-resident chaperone, heat shock protein 47 (HSP47) along with other ER foldases such as BIP/GRP78 (4, 5). Under normal physiological conditions, HSP47 is expressed at low levels in the liver (6) and other organs such as lung, heart, kidney (7). Fibrogenic stimulation by carbon tetrachloride (CCl4) or bile duct ligation (BDL) in mice and human liver fibrosis is associated with induction of HSP47 expression (6,8,9). The induction in HSP47 correlates with increased collagen secretion from activated HSCs during liver fibrosis. Therefore, silencing HSP47 to inhibit collagen production is an appealing option for reversing fibrosis (10). However, because HSP47 also plays a chaperoning function in the healthy liver and other organs, the collateral effects of its therapeutic silencing should be investigated (10). Apart from its function as a collagen chaperone, a recent interactome study identified HSP47 as a binding partner for an unfolded protein response (UPR) sensor protein, inositol-requiring enzyme 1 alpha (IRE1α) (5). HSP47 activates IRE1α oligomerization and phosphorylation by displacing its regulator, BIP, thereby triggering the UPR response during ER stress (5). Whether triggering of UPR signaling by HSP47-IRE1α interaction and BIP displacement in HSCs may enhance folding of pro-fibrogenic proteins such as collagen is unclear so far. But it is generally accepted that HSCs exhibit ER stress and UPR signaling in response to liver injury stimuli (11).

A-kinase anchoring protein 12 (AKAP12) is a ubiquitously expressed member of the AKAP family that exhibits scaffolding activity towards signaling molecules including protein kinases (PKA and PKC), β2-adrenergic receptor, cyclins-(cyclin-D1, CCND1) (12) and polo-like kinase 1 (PLK1) (13). By virtue of its scaffolding function, AKAP12 spatiotemporally controls cellular signaling by guiding its binding partners to their physiological substrates or specific functional locations (14). These activities regulate growth, cytoskeletal remodeling and adrenergic signal transduction (12). AKAP12-mediated scaffolding of PKC attenuates PKC activation and suppress oncogenic proliferation, invasiveness, chemotaxis, and senescence (15, 16). PKCα, δ and ε isoforms interact with AKAP12, however only PKCα and δ activity is induced in the absence of AKAP12 (17, 18). AKAP12 sequestration of CCND1 in the cytoplasm prevents its nuclear translocation and cell cycle progression (19). AKAP12 sequestering of CCND1 and inhibition of CCND1 activity has been reported in parietal glomerular epithelial cells and in fibrosarcoma (20, 21).

It has been demonstrated that the scaffolding ability of AKAP12 is altered by its phosphorylation (17, 19, 22). Prephosphorylation of AKAP12 by PKC suppresses its interaction with PKC itself and increases PKC activity (23). Phosphorylation of AKAP12 at a PKC phosphorylation site (S507/515) prevents the sequestration of CCND1 by AKAP12 leading to its nuclear translocation, allowing cell cycle progression (19, 21). AKAP12 phosphorylation by cyclin-dependent kinase 1 (CDK1) at a threonine residue (T766) enhances the recruitment of the polo-like kinase (PLK1) in human glioblastomas to ensure efficient mitotic progression (13). Even though phosphorylation is known to regulate AKAP12’s scaffolding activities, the functional impact of its phospho-modifications on liver disease has not been evaluated. We previously demonstrated that HSC activation during liver injury was associated with an induction in phospho-AKAP12 (24). In this work we demonstrate that specific AKAP12 phosphorylation events in HSCs regulate its scaffolding activity towards the collagen chaperone, HSP47. HSC-specific CRISPR-editing of AKAP12’s phospho-sites preserves the AKAP12-HSP47 scaffold, reduces HSP47’s collagen-chaperoning activity, dramatically lowering overall collagen content and liver injury during carbon-tetrachloride (CCl4)-induced liver fibrosis. AKAP12 phospho-modulation directed towards HSCs regulates HSP47-IRE1α interaction, thereby controlling UPR signaling in HSCs. Furthermore, AKAP12 phospho-site modulation in HSCs suppresses overall ER stress in the fibrotic liver. Our data supports a previously unidentified function of AKAP12 and its phospho-modification in regulating the outcome of liver fibrosis in animal models.

## Results

### Expression, phosphorylation, and scaffolding activity of AKAP12 is altered in CCl4-treated mouse liver and human liver fibrosis

The expression of AKAP12 protein was decreased in livers of CCl4-treated mice by 14% compared to oil controls without a change in *Akap12* mRNA (Figure 1A). CCl4 treatment induced the expression of HSC activation marker, α-SMA by 4.5-fold compared to control (Figure 1A, figure 1-source data 1). As evidenced by proximity ligation assay (PLA), the phosphorylation of AKAP12 was induced in desmin-positive HSCs of CCl4 livers by 5.3-fold compared to control (Figure 1B, figure 1-source data 2). AKAP12 staining judged by Image J quantification (materials and methods) was decreased in CCl4-treated liver by 16% compared to control (Figure 1B, figure 1 source data 2) consistent with the western blot result (Figure 1A). The interaction of AKAP12 with HSP47 was inhibited by 54% despite a 3.9-fold increase in overall HSP47 levels in CCl4 livers compared to control (Figure 1C, figure 1-source data 3). A human liver fibrosis tissue array containing 16 liver fibrosis tissues and 11 normal tissues was stained with PLA probes for AKAP12 and HSP47 to detect their interaction. The interaction between AKAP12 and HSP47 was inhibited by 64% in human liver fibrosis tissue compared to normal (Figure 1D, figure 1-figure supplement 1). This was associated with a 20% decrease in total AKAP12 staining and a 3.8-fold increase in HSP47 staining in liver fibrosis compared to normal (Figure 1D).

**Figure 1.**
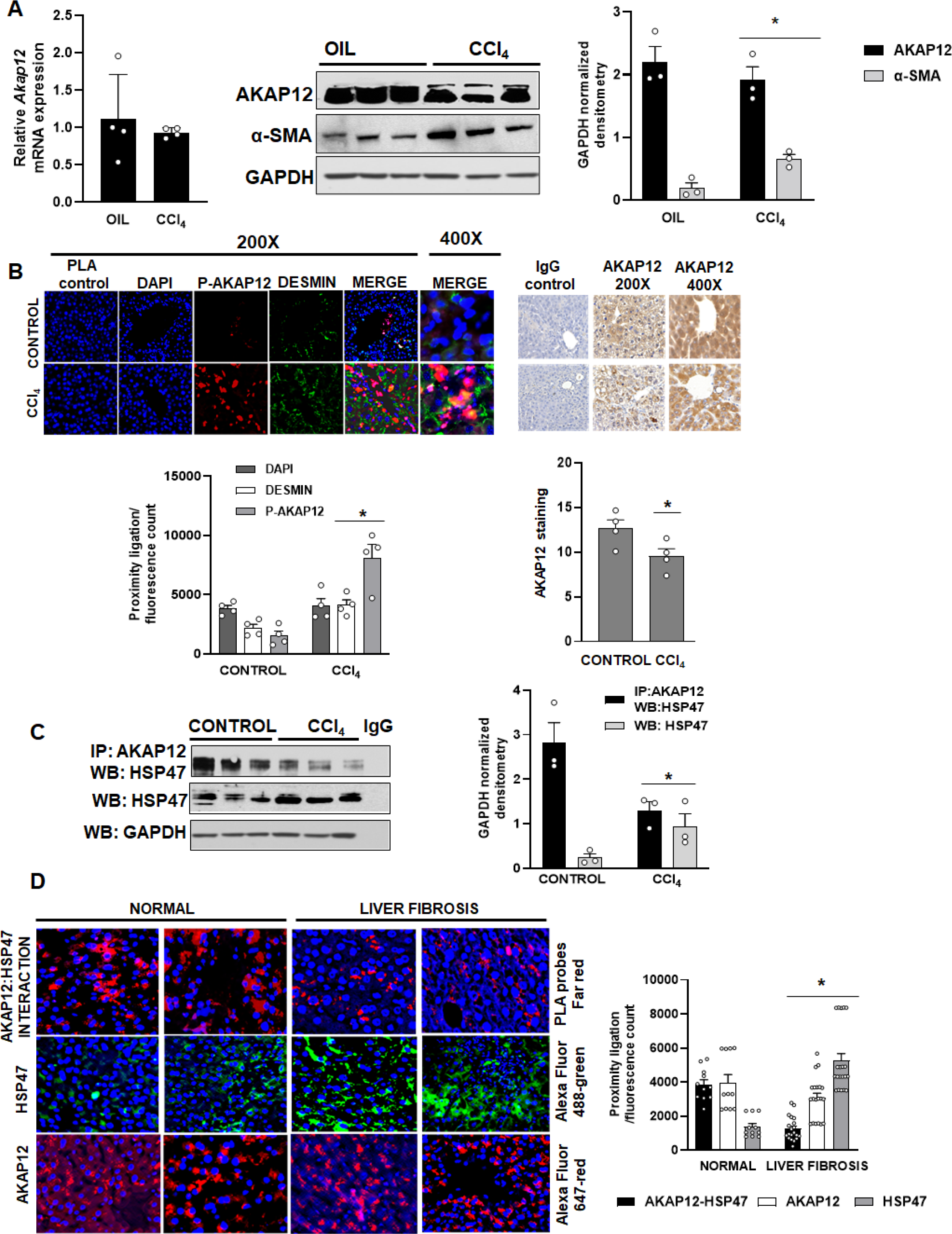
Expression, phosphorylation, and scaffolding activity of AKAP12 is altered in CCI_i_-treated mouse liver and human liver fibrosis. Mice were administered CCi_4_ or mineral oil (control) as in methods. (A) Total RNA (left panel) from mouse liver was subjected to real-time RT-PCR to evaluate the expression of Arapl2or Gapah (normalizing control) mRNA (Mean±S.E from 4 experimental groups). Total protein (right panel) was immuπob lotted with AKAP12_1_ α-SMA or GAPDH (control) antibody and blots were quantified by Image J densitometry’. Data represented by GAPDH normalized densitometry’ is mean±S. E from 3 experimental groups. AKAP12: ’p=0.Q02_t_ a-SMA: *p=0.04 vs. control. (B) Sections of control or CC∣_1_ livers stained with the HSC marker, desmin was overlayed with antibodies to detect phospho-AKAP12 (P-AKAP12) by PLA as in methods. Total AKAP12 expression was detected by HRP/DAB staining as in methods. Images were quantified by densitometry using Image J and represented as the proximity ligation/fiuorescence/HRP count. Mean±S.E from4 experimental groups. Desmin: *p=0.012_t_ P-AKAP12 PLA: *p=0.0017_l_ AKAP12: ^i^p=D.D4 vs. control. (C) Control or CCi_4_ liver protein was immu∩oprecipitated with AKAP12 antibody and probed for HSP47 by western blotting. Normal mouse IgG was a negative control. Data represented by GAPDH normalized densitometry are mean±S.E from 3 experimental groups. AKAP12-HSP47 Co-IP: “p=□.□014. HSP47: *p=0.004 vs. control. (D) Human tissue arrays were stained with AKAP12 and HSP47 far red PLA probes as in methods. AlexaFluor antibodies (supplementary’ table S6) were used to detect expression of AKAP12 or HSP47 in these arrays. A representative area is shown at 400X magnification. The complete array staining at 100X magnification is shown in supplementary figure 1. Each tissue within the array was quantified by densitometry using the Image J and represented as the proximity ligatio n/fluorescenee count. Mean±S.E, from 11 normal livers and 16 liver fibrosis tissues. AKAP12- HSP47 PLA: *p=0.013, AKAP12 staining: *p=0.03E, HSP47 staining: *p= 1.98X10-6 vs. normal.

### CRISPR-directed editing of AKAP12’s activation-responsive phospho-sites enhances AKAP12’s scaffolding activity and inhibits HSC activation

The phospho-peptide map of AKAP12 protein from Day 7 culture-activated human or mouse HSCs was compared to that of Day 0 quiescent HSCs or normal hepatocytes. A peptide region containing 5 S/T phospho-sites exhibited increased phosphorylation in Day 7 activated HSCs but not in Day 0 HSCs or hepatocytes (Table 1, table supplement 1). These activation-responsive phospho-sites were conserved in mouse and human (Table 1). Day 5 activated human HSCs were transfected with CRISPR small guide RNA (sgRNA) and donor RNA (Supplementary table S1) to delete the 5 AKAP12 phosphorylation sites by homology-directed repair (HDR) as described under materials and methods. Genomic DNA PCR from CRISPR edited (HDR) cells using deletion-specific primers (Supplementary table S1) resulted in a 261 bp amplicon that was not amplified in WT cells or cells treated with SaCas9 (staphylococcus aureus CRISPR-associated protein) alone (Figure 2A, original gel shows 4 experiments). The interaction between AKAP12 and HSP47 in CRISPR-edited HSCs (HDR) was induced by 2.5-fold compared to WT cells (Figure 2B, original blot developed with anti-mouse IgG is shown in figure 2-source data 1). This was associated with a 40% decrease in α-SMA levels, demonstrating that AKAP12 phospho-site editing inhibited HSC activation (Figure 2B). The overall level of HSP47 decreased by 25% whereas the level of AKAP12 protein remained unchanged after HDR (Figure 2B). Deletion of phospho-sites in mouse HSCs resulted in a 422 bp deletion-specific amplicon (Figure 2-figure supplement 1-source data 1). Like human HSCs, mouse HSCs also exhibited increased AKAP12-HSP47 interaction after AKAP12 phospho-site editing (figure 2-figure supplement 1- source data 2). Reversal of HSC activation by AKAP12 editing was determined by examining vitamin A auto fluorescence (25). Cultured human HSCs at Day 0 exhibited strong vitamin A autofluorescence that was reduced in Day 5 activated HSCs (Figure 2C, 3 independent experiments are shown). AKAP12 editing in Day 5 HSCs restored the loss of vitamin A fluorescence compared to Day 5 HSCs or Day 5 HSCs+SaCas9 alone (Figure 2C). HSP47 is an ER-resident chaperone (4). A weak PLA signal of AKAP12-HSP47 interaction co-localized with the ER marker, calreticulin in activated (WT) HSCs (Figure 2D, left panel). However, upon CRISPR-editing (HDR) a strong AKAP12-HSP47 PLA signal co-localized with calreticulin in the ER (Figure 2D, left panel). We examined whether HSP47’s collagen-chaperoning activity was regulated by AKAP12 phospho-site editing. Our results show that the collagen-HSP47 PLA signal strongly co-localized in the ER of activated HSCs (WT) (Figure 2D, right panel). CRISPR-editing of AKAP12 (HDR) reduced the collagen-HSP47 interaction significantly by 65% compared to WT cells (Figure 2D, right panel). Individual experiments are shown in figure 2-source data 2.

**Figure 2.**
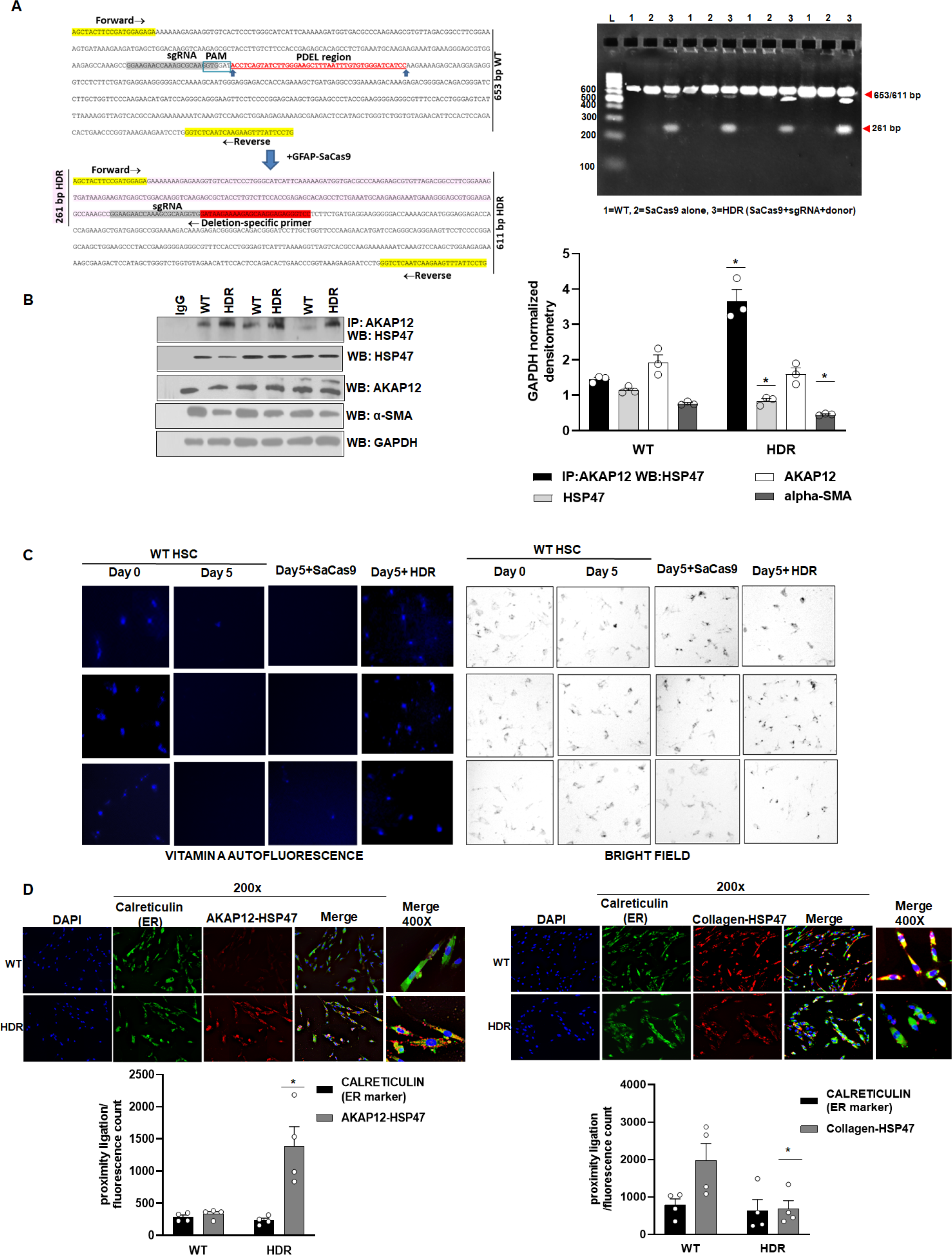
CRISPR-directed editing of AKAP12’s activation-responsive phospho-sites enhances AKAP12’s scaffolding activity and inhbits HSC activation. Actuated human HSCs were transfected with CRISPR reagents and GFAP-SaCas9 vector to cause CRIS PR-directed HDR as in methods. Untransfected (WT) or cells with SaCas9 alone were used as controls. (A) CRISPR editing at the AKAP12 locus (left panel) was confirmed by performing PCR (right panel) using primers that specifically detected the edited region as listed in supplemental table S1. Four independent experiments are shown. (B) CRISPR-edited (HDR) or WT cells were assessed for AKAP12-HSP47 co-immunoprecipitation, HSP47, AKAP12 and α-SMA (HSC activation marker) western blotting. Data represented as GAPDH normalized densitometry is mean±S.E from 3 experiments. AKAP12-HSP47 Co-IP: *p=0.009, HSP47: *p=0.03, a-SMA: *p=0.0002 vs. WT cells. (C) Day 0 attached HSCs were culture activated till Day 3 and then transfected with CRISPR vectors till Day 5. The autofluorescence of vitamin A as a marker of HSC quiescence was visualized by fluorescence microscopy and compared to brightfield images of cells as in methods. Three independent experiments are shown. (D) AKAP12-HSP47 interaction (left panel) and HSP47-collagen interaction (right panel) in the ER was compared between WT and HDR cells by PLA staining and co-staining with the ER marker, calreticulin as in methods. Data represented as proximity ligation/fluorescence count are mean±S.E from 4 experiments. AKAP12-HSP47 PLA: *p=0.014, Collagen-HSP47 PLA: *p=0.04 vs. WT cells.

**Figure 3.**
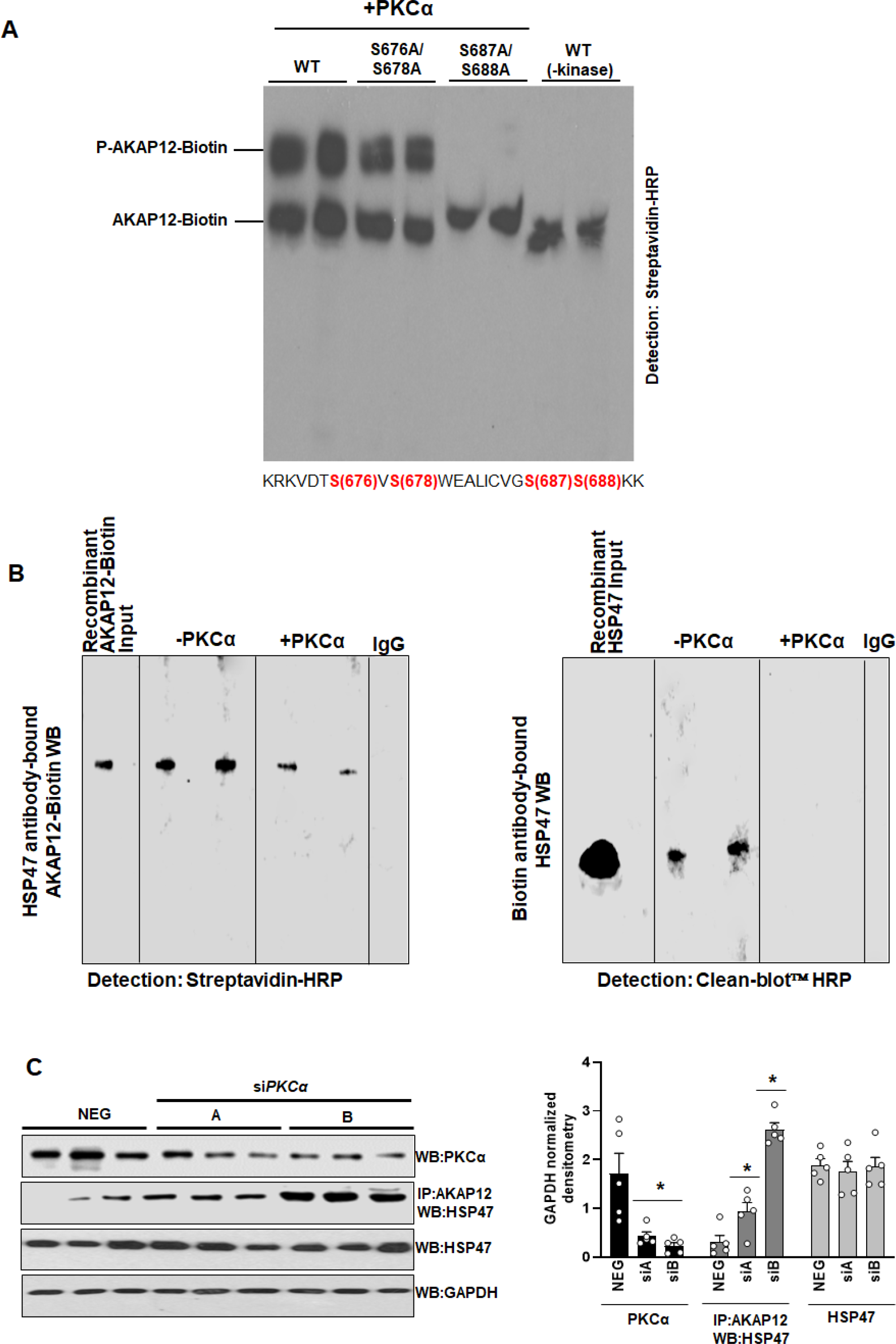
PKCa phosphorylâtes AKAP12 and inhibits its interaction with HSP47. (A) AKAP12 is phosphorylated by PKCα at its activation-responsive phospho-sites. Recombinant WT or AKAP12 phospho-mutants were in vitro translated from their vectors and subjected to in vitro kinase assay in the presence of active PKCo enzyme as in methods. The reactions were run on a phostag^r^* gel to detect phosphorylated AKAP12 or its mutants. Representative phostag∼ gels from 3 experiments are shown. (B) Direct Interaction between AKAP12 and HSP47 in recombinant system in the absence or presence of active PKCσ enzyme. In vitro translated biotinylated AKAP12 was incubated with recombinant HSP47 antibody column containing bound HSP47 (left) or recombinant HSP47 was incubated with Biotin antibody column containing bound AKAP12-Biotin (right) in the presence or absence of active PKCα as in methods. Two representative data out of 4 experiments are shown. (C) Silencing PKCa in activated human HSCs enhances AKAP12-HSP47 interaction. Culture-activated human HSCs were transfected with a universal negative control siRNA (Neg) or two PKCa siRNAs (A or B) as in methods. Total protein was assessed for AKAP12-HSP47 co-immunoprecipitation or PKCa, HSP47 and GAPDH immunoblotting. Data represented asGAPDH normalized densitometry is mean±S.E from 5 experiments. PKCa: *p=0.002, AKAP12-HSP47 Co-IP: *p=0.04 vs. Neg.

**Table 1.** Phospho-peptide mapping of human HSCs, mouse HSCs and mouse hepatocytes

| Cell type | Observed precursor mass | Neutral Loss of Phosphate Mass | Phospho-Peptide Sequence | Peptide modification |
| --- | --- | --- | --- | --- |
| Day 0 human HSC | 1988.7812 | 1890.0297 | KRKVDTSVSWEALICVGSSKK | Phospho (ST)[16] |
| Day 7 human HSC | 2148.9758 | 1854.9 | KRKVDTSVSWEALICVGSSK | Phospho (ST)[16,17], |
| Day 7 human HSC | 2148.9932 | 1855.3141,<br>1854.9 | KRKVDTSVSWEALICVGSSKK | Phospho (ST)[4,6,16] |
| Day 0 mouse HSC | ND | ND | KRKVDTSVSWEALICVGSSKK | ND |
| Day 7 mouse HSC | 1998.13 | 1801.7952 | KRKVDTSVSWEALICVGSSK | Phospho (ST)[3,6] |
| Day 7 mouse HSC | 2054.22 | 1857.9027 | KRKVDTSVSWEALICVGSSKK | Phospho (ST)[14,15] |
| mouse hepatocytes | ND | ND | KRKVDTSVSWEALICVGSSK | ND |

S=Serine, T=Threonine, ND=not detected

### PKCα phosphorylates AKAP12 and inhibits its interaction with HSP47

Kinase-prediction software were used to predict that out of the 5 AKAP12 activation-responsive phospho-sites, two serines (S687/S688) were strongly predicted substrates of PKCα kinase with a consensus of [S/T]-X-R/K whereas one threonine (T675) could not be assigned a kinase (supplementary table S2). S676/S678 sites were also PKCα sites but shared consensus sites with calmodulin kinase (CAMK). The overall confidence of prediction for the S676/S678 sites was less than that of S687/S688 sites. In vitro kinase assay followed by phostag™ gel analysis revealed that phosphorylation of AKAP12 was significantly enhanced in the presence of active PKCα enzyme compared to kinase negative controls (Figure 3A). Mutations of AKAP12 S676/S678 to alanine modestly reduced the phosphorylation of biotinylated recombinant AKAP12 whereas S687A/S688A mutation dramatically suppressed the phostag™ shift of AKAP12 (Figure 3A). The mutation seemed to completely inhibit the phospho-band. Since other phosphorylation events could also cause the shift, we repeated the experiment to see whether this complete suppression was reproducible. In an additional experiment (Figure 3-source data 1) we observed that the S687/S688A mutation suppressed but did not always wipe out the phospho-shift. Also, in some experiments we observed the -kinase control had a faint phospho-signal. The recombinant protein produced by rabbit reticulocyte lysates in an in vitro translated system may have baseline phosphorylation as reported in the manufacturer’s protocol (TNT® Coupled Transcription/Translation system, Promega). Direct binding was observed between biotinylated AKAP12 and HSP47 in a recombinant system in the absence of active PKCα (Figure 3B, figure 3- source data 2). Presence of PKCα inhibited the interaction between AKAP12 and HSP47 (Figure 3B). To evaluate whether phosphorylation of AKAP12 by PKCα in HSCs would regulate AKAP12’s scaffolding activity, cells were treated with *PKCα* siRNAs (A or B). Silencing *PKCα* by 74% with siRNA-A increased AKAP12-HSP47 interaction by 3-fold whereas a 90% knockdown caused by siRNA-B enhanced AKAP12-HSP47 interaction by 8-fold compared to negative control siRNA (Figure 3C, figure 3-source data 3). HSP47 levels remain unchanged by siRNA treatments (Figure 3C).

### In vivo gene editing of the *Akap12* region corresponding to its activation-responsive phospho-sites in HSCs of mouse liver

The *Akap12* exon 3 contains sequences corresponding to the activation-responsive phospho-sites of AKAP12 protein. To perform gene editing of this region specifically in HSCs of mouse liver, two different CRISPR HDR approaches were used (Figure 4A). A PDEL donor was used to delete the AKAP12 phospho-sites whereas each S or T phospho-site was mutated to A using a PMUT donor. Two unique sgRNAs specific for the region around the phospho-sites along with the donor (Figure 4A, Supplementary table S1) were cloned into AAV vectors (Figure 4B, left panel). To perform CRISPR editing in HSCs of mouse liver, the SaCas9 enzyme was cloned into AAV vector under control of two different HSC-specific promoters (Glial fibrillary acidic protein, GFAP or Lecithin retinol acyltransferase, LRAT) (26, 27). AAV vectors were injected into mice during oil or CCl4 administration according to the plan in Figure 4B, right panel. To evaluate the HSC specificity of GFAP-SaCas9 mediated CRISPR (CR) editing compared to that of an empty vector (EV) control (materials and methods), genomic DNA of HSCs or hepatocytes isolated from livers of oil+EV, oil+CR, CCl4+EV, CCl4+CR groups was subjected to multiplex PCR with PDEL forward and reverse primers and a PDEL deletion-specific primer (Supplementary table S1). Oil+EV or CCl4+EV HSCs or hepatocytes gave a 298 bp amplicon in this multiplex PCR. Oil+CR or CCl4+CR groups resulted in 298 bp WT and 256 and 154 bp mutated amplicons due to complementarity with the deletion-specific primer (Figure 4C). PCR with deletion-specific primers did not amplify the 256 or 154 bp mutant region in hepatocytes, indicating that CRISPR-editing using an HSC promoter-specific SaCas9 occurred in HSCs but not hepatocytes (Figure 4E). HSC specificity of CRISPR was also evaluated by immunostaining of Sacas9 enzyme with HSC marker, desmin or hepatocyte marker, albumin. The GFAP-driven SaCas9 enzyme strongly co-localized with desmin-positive HSCs (Figure 4D) but not with albumin-positive hepatocytes in the liver (Figure 4F). CCl4 exposure increased the overall numbers of desmin-positive HSCs (Figure 4D) due to increased activation and proliferation (28). A specific primer to detect PMUT could not be designed, hence PMUT specificity and efficiency was tested along with PDEL by next generation amplicon sequencing (NGS) using a 298 bp amplicon from HSCs or hepatocytes of GFAP-SaCas9 CRISPR livers (Figure 4G). On-target and off-target base changes were analyzed by comparing the target read sequences to the reference sequence of WT *Akap12* amplicon as described under materials and methods. For the PDEL CRISPR, Oil+CR HSCs exhibited 30% mutant reads compared to the total reads whereas CCl4+CR HSCs exhibited 60% mutant reads compared to total (Figure 4G, top panel, supplementary table S3). Oil+EV or CCl4+EV HSCs did not contain any mutant reads (Figure 4G, top panel). For the PMUT CRISPR, oil+CR HSC exhibited 3% mutant reads and CCl4+CR exhibited 12.5% mutant reads compared to total reads (Figure 4G, botom panel, supplementary table S4). Hepatocytes from the CR groups did not exhibit any PDEL or PMUT sequence reads (Figure 4G, supplementary tables S3 and S4). The percentage of base changes outside the target region between the two sgRNA sites (Figure 4A) was less than 5% in most cases (Figure 4G). Like GFAP-SaCas9, HSC-specific PDEL CRISPR was also observed with LRAT-SaCas9 (figure 4-figure supplement 1). The LRAT-driven SaCas9 expression in desmin-positive HSCs was lower in the CCl4 group compared to oil (figure 4-figure supplement 1). The CRISPR deletion efficiency using LRAT-saCas9 in HSCs of oil+CR group was 45% whereas that of the CCL4+CR group was 30% of total reads (figure 4-figure supplement 1, Supplementary table S3).

**Figure 4.**
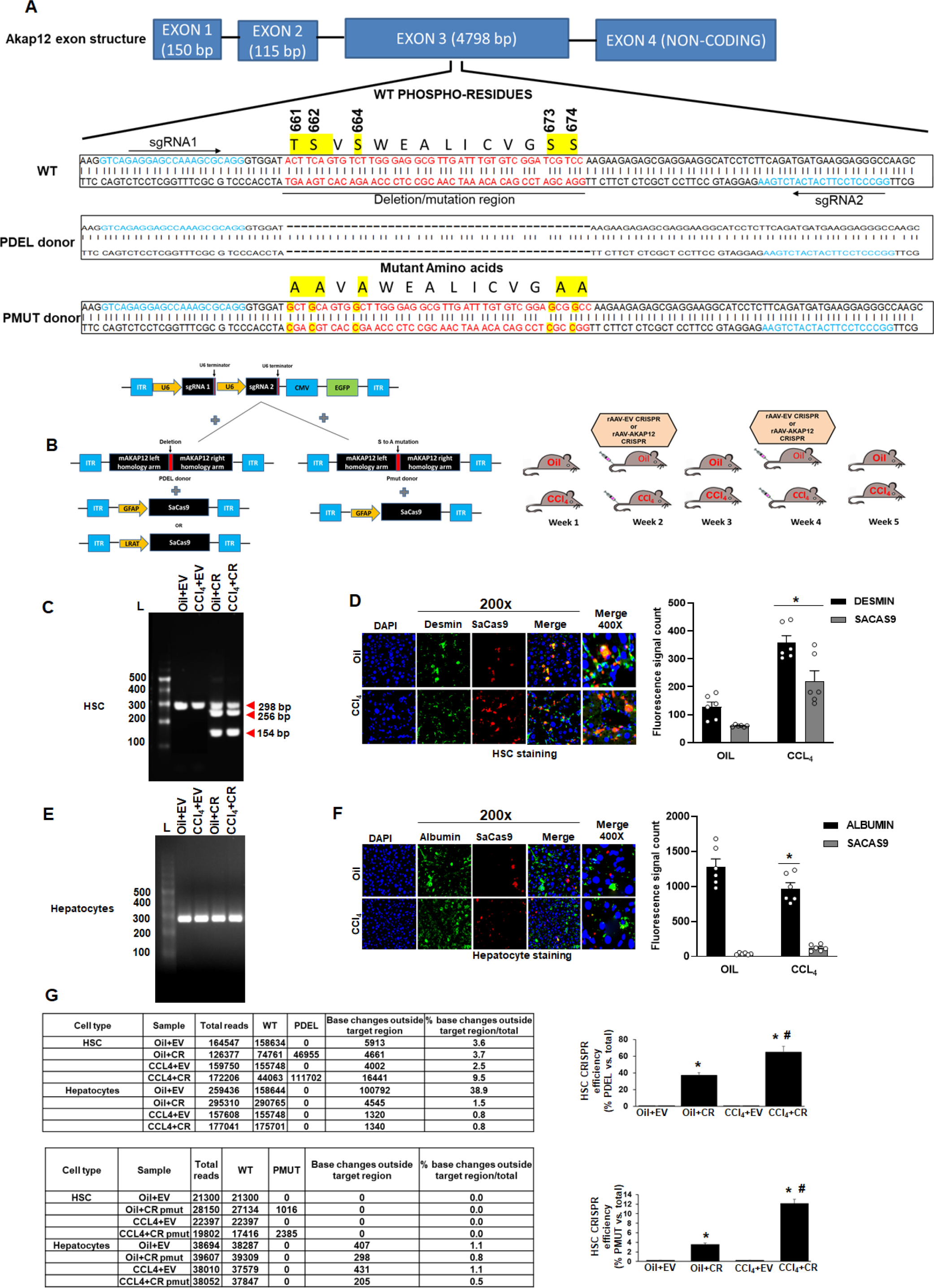
In vivo gene editing of the Akap12 region corresponding to its activation-responsive phospho-sites in the CCL4 mouse model using GFAP-SaCas9. (A) Schematic diagram of the mouse Akap12 locus showing the exon 3 region containing AKAP12’s activation-responsive phospho-site regions and two SaCas9 target sgRNAs (1 and 2). The PDEL mutation contains a 42-bp deletion in the donor that deletes the phospho-sites after CRISPR editing. The PMUT donor has a mutation in 5 codons that changes the S/T (serine/threonine) activation-responsMe phosphorites to A (alanine). (B) Left panel: AAV-CRISPR cloning scheme. SgRNA1∕2, PDEL or PMUT donor and SaCas9 under control of the HSC-specific GFAP promoter were cloned into AAV6 serotype vectors and AAV particles were generated as in methods. Control vector (EV) contained a non-targeting sgRNA as in methods. Combinations of sgRNA1∕2-AAV or EV with the pDEL or pMUT donor-AAV and GFAP-SaCas9-AAV were injected into mice as in methods. Right panel: Scheme of AAV vector injections into the tail vein at 2nd and 4th week of oil or CCL4 administration as in methods. (C) Specificity of PDEL CRISPR for HSCs was evaluated by multiplex PCR amplification of genomic DNA from HSCs using a PDEL-specific primer and two primers around the PDEL primer site (supplementary table 1). A representatize gel image from 6 experimental groups is shown. (D) Specificity of HSC CRISPR was ascertained by co-localization of SaCas9 with desmin-positive HSCs in oil or CCL4-treated groups. Data represented by fluorescence signal count are mean±S.E from 6 experimental groups. Desmin staining: *p=0.011, desmin-positive SaCas9 staining: *p=0.04 vs. Oil. (E) Multiplex PCR of hepatocytes genomic DNA as in C above did not show PDEL specific amplicons. A representative gel image from 6 experiments is shown. (F) SaCas9 co-localization with albumin-positive hepatocytes was insignificant in oil or CCL4 livers compared to HSCs in D’ above. Data represented by fluorescence signal count are mean±S.E from 6 experimental groups. Albumin staining: *p=0.05 vs. oil+EV. (G). The efficiency of CRISR was evaluated NGS using a 298 bp PCR amplicon derived from genomic DNA of HSCs or hepatocytes of PDEL mice group (top panel) or PMUT mice group (bottom panel). Total amplicon reads, WT reads, and PDEL or PMUT reads within the target region or base changes outside the target region from each experimental group are shown. The CRISPR editing efficiency represented by the percentage of mutant reads versus total is mean±S.E from6PDEL or 3 PMUT experimental groups. Oil+CR orCCL4+CR-PDEUPMUΓ *p=0.0002 vs. Oil+EV; CCL4+CR PDEL/PMUT: #p=0.0003 vs. CCL4+EV.

### Phospho-editing of AKAP12 regulates liver injury and fibrosis in the CCl4 mouse model

At gross level, CCl4 administration for five weeks reduced the body weight of mice by 20% and increased the liver to body weight ratio by 1.25-fold compared to oil (Figure 5A). AKAP12 phospho-editing by GFAP-SaCas9 in normal mice (oil+CR) did not alter body or liver weight compared to oil+EV (Figure 5A). However, AKAP12 phospho-editing in CCL4 mice (CCl4+CR) normalized CCL4+EV-mediated alterations in body weight and liver/body weight to that of oil+EV levels (Figure 5A). Histologically, control mice (oil) had a normal hepatic cord pattern around the central vein whereas fatty vacuolar changes and disorganized hepatic lobular structure with centrilobular fibrosis was observed in CCl4 livers (Figure 5B) as referenced previously (29). This was associated with an 8-13-fold induction in liver injury as measured by ALT/AST levels (Figure 5C). AKAP12 phospho-editing by PDEL or PMUT in control mice (oil+CR) did not affect the normal histology of the liver or the levels of ALT/AST (Figure 5B, C). AKAP12 phospho-editing by PDEL or PMUT in CCl4 mice (CCl4+CR-PDEL or PMUT) dramatically reduced the CCl4-induced histological distortions and decreased the ALT/AST level by 75-80% compared to CCl4+EV (Figure 5B, C). H&E staining for 6 PDEL experiments is shown in figure 5-source data 1. LRAT-SaCas9 directed PDEL-CRISPR also resulted in higher body weight, lower liver/body weight ratio and suppression of CCl4-induced histological changes like that of GFAP-SaCas9 (Figure 5-figure supplement 1).

**Figure 5.**
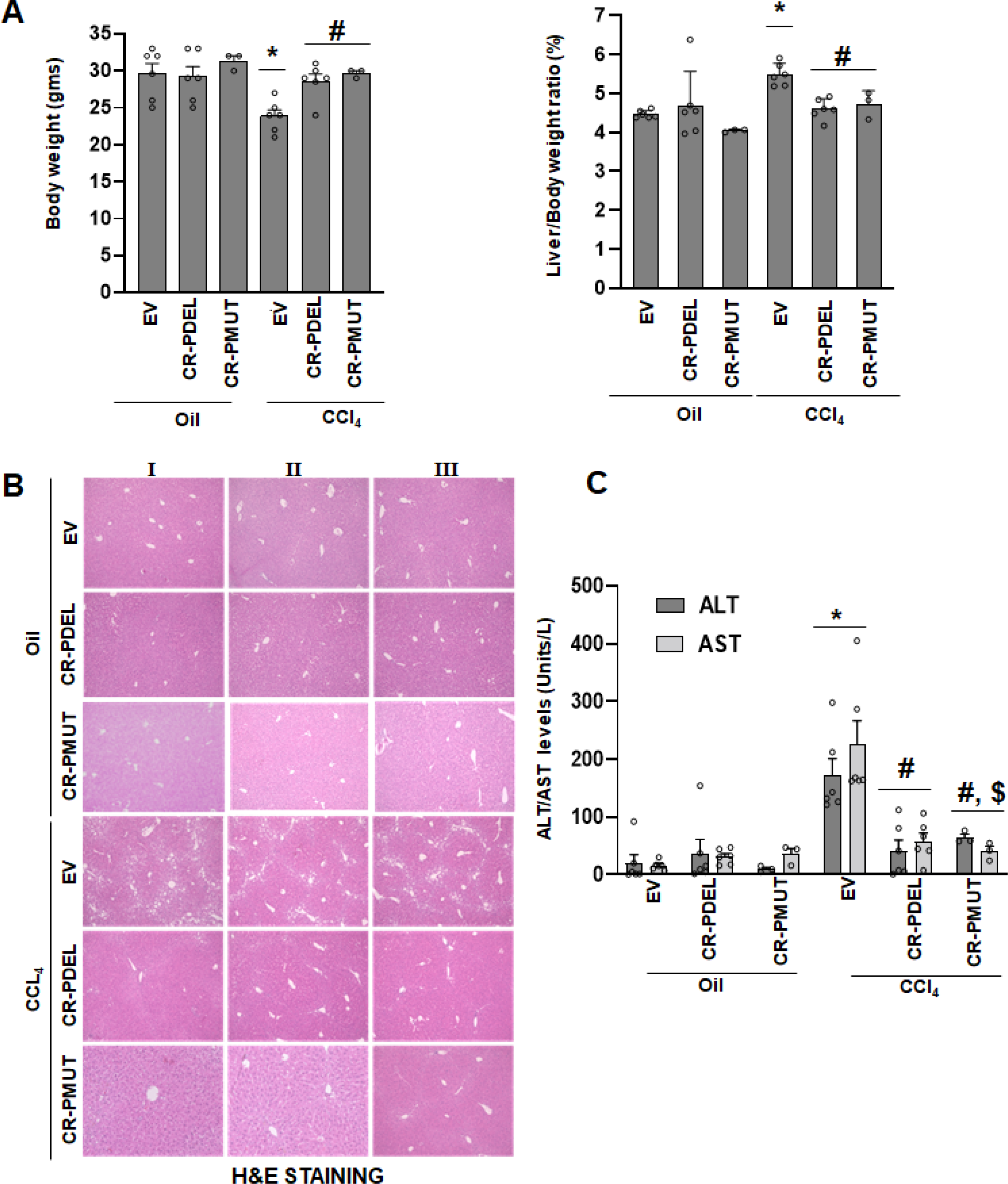
Ptospho-editing of AKAP12 regulates liver injury and fibrosis in the CCL_4_ mouse model. (A) Gross changes In mouse body weight (left panel) and liver/body weight ratio [right panel) after PDEL or PMUT GFAP-SaCasE*-mediated CRISPR editing of AKAP12’s phospho-sites under oil or CCL_j_ treatment conditions. Mean ±S.E from 5 PDEL experiments and 3 PMUT experiments. CCU_4_+EV: *p=0.0001 vs. oil+EV; CCL_4_+CR-PDEL <0=0.008, CCL_4_+CR-PMUT: #p=0.Q2 vs. CCL_4_+EV. (B) Histological evaluation of CRISPR-edited livers by H&E staining as In methods. Three cut of 6 PDEL experiments and 3 PMUT experiments are shown. [C) Measurement of ALT and AST levels In plasma of after CRISPR-editing is In methods. Mean±S.E from 5 PDEL experiments and 3 PMUT experiments. CCL_4_+EV *P=0.008 vs. oll+EV; CCL_4_+CR-PIDEL *p=0.02, CCL_4_+CR-PMUT #p=0.0009 vs. CCL_4_+EV; CCL_4_+CR-PMUT $p=0.004 vs. oil+EV or oil+CR.

### Phospho-editing of AKAP12 regulates AKAP12’s HSP47-scaffolding activity, HSC activation, HSP47’s collagen-chaperoning activity and collagen production in the CCl4 mouse model

The AKAP12-HSP47 scaffold was reduced in livers of CCl4+EV mice compared to oil controls (Figure 6A and B). PDEL-CRISPR or PMUT-CRISPR editing in CCl4 mice restored the drop in the AKAP12-HSP47 interaction caused by CCl4 (Figure 6A, PDEL; figure 6B, PMUT, figure 6-source data 1, PDEL; figure 6-source data 2, PMUT). AKAP12 phospho-editing dramatically inhibited CCl4-mediated HSC activation as evidenced by a drop in α-SMA levels (Figure 6A). In conjunction with restoration of AKAP12-HSP47 scaffold, the increased interaction between collagen and HSP47 upon CCl4 exposure was inhibited by AKAP12 phospho-editing (Figure 6C). AKAP12 PDEL or PMUT phospho-editing also inhibited the increase in collagen mRNA levels caused by CCl4 exposure (Figure 6C). Co-immunoprecipitation of collagen with HSP47 antibody yielded non-specific bands at positions above the collagen position in all samples including IgG control. The original uncropped blot is shown in figure 6-source data 3. PLA staining showed that the AKAP12-HSP47 scaffold was localized with desmin-positive HSCs under normal (oil) conditions (Figure 6D). A drop in AKAP12-HSP47-desmin co-localization was observed upon CCl4 exposure that was restored by AKAP12 PDEL or PMUT phospho-editing (Figure 6D). Picosirus red staining of CCl4 livers showed increased collagen deposition that was substantially reduced when mice were administered AKAP12 phospho- editing vectors (Figure 6E, top panel). Sirius red staining for 6 PDEL experiments is shown in figure 6-source data 4. The hydroxyproline content of collagen was increased 2.4-fold in CCl4 livers compared to oil+EV and normalized by AKAP12 PDEL. PMUT phospho-editing inhibited CCl4-mediated induction but did not completely normalize hydroxyproline content to oil+EV or oil+CR levels (Figure 6E, bottom panel).

**Figure 6.**
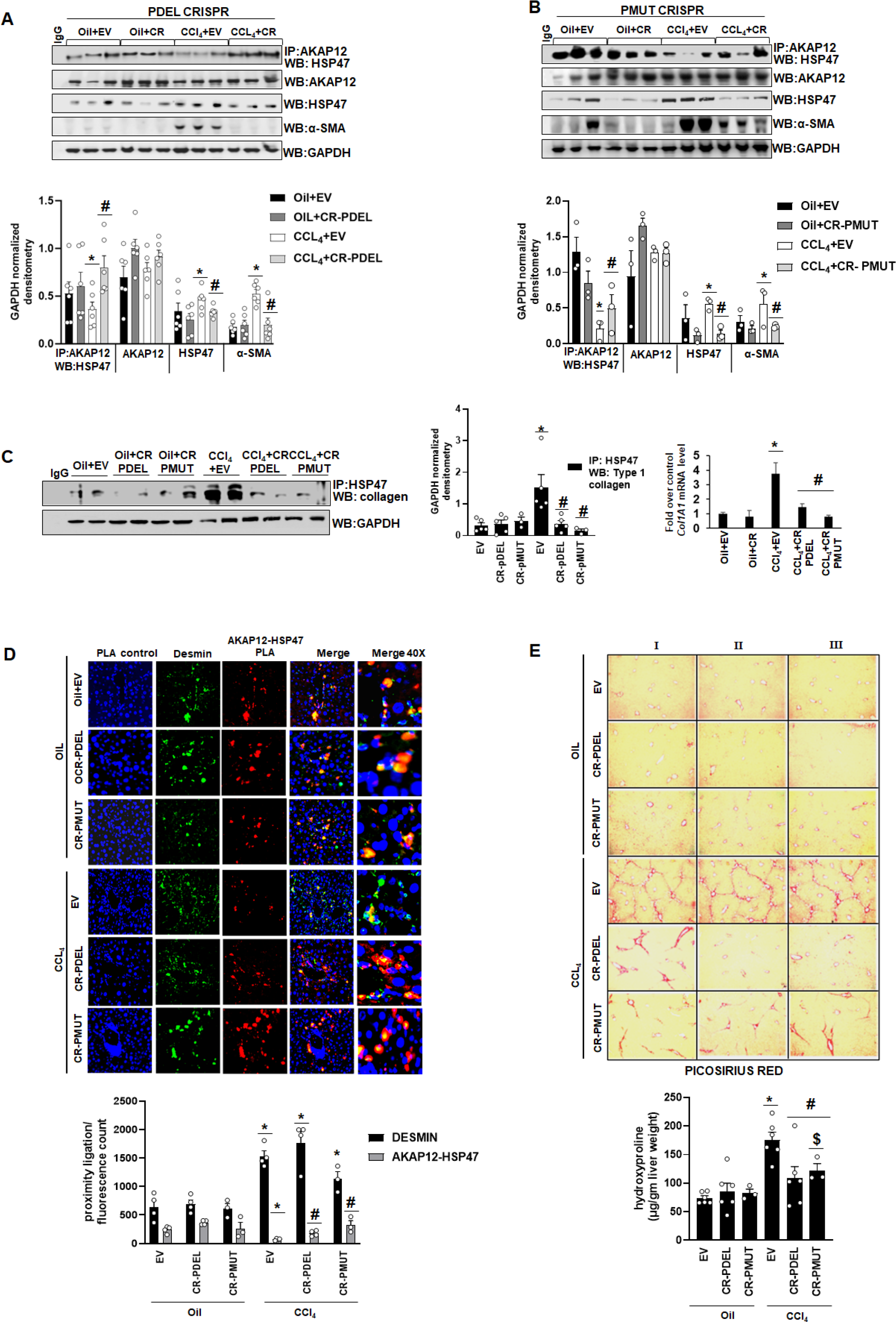
Phospho-editing of AKAP12 regulates AKAP12’s HSP47-scaffolding activity, HSC activation, HSP47’s collagen-chaperoning activity and collagen production in the CCI4 mouse model. (A). AKAP12-HSP47 co-immunoprecipitation, AKAP12, HSP47 and α-SMA western blotting from liver protein of CR-PDEL experiment. Data represented as GAPDH normalized densitometry is mean±S.E from 6 experiments. Three representatives are shown. AKAP12-HSP47 Co-IP: *p=0.04, HSP47: *p=0.001, a-SMA: *p=0.009 vs. oil+EV; AKAP12-HSP47 Co-IP: #p=0.018, HSP47: #p=0.04, a-SMA: ⅛p=0.003vs. CCL4+EV. (B) AKAP12-HSP47 co-immunoprecipitation, AKAP12, HSP47 and α-SMA western blotting from liver protein of CR-PMUT experiment. Data represented as GAPDH normalized densitometry are mean+S.E from 3 experiments. All three representatives are shown. AKAP12-HSP47 Co-IP: *p=0.008, HSP47: *p=0.016, a-SMA: *p=0.004 vs. oil+EV. AKAP12-HSP47 Co-IP: ⅛p=0.049, HSP47: #p=0.004, a-SMA: #p=0.0002 vs. CCL4+EV. (C) Left panel: Co-immunoprecipitation of collagen with HSP47 in CRISPR-edited livers, *p=0.019vs. oil+EV; CCI4+CR-PDEL: #p=0.02, CCI4+CR-PMUT: #p=0.009 vs. CCL4+EV. Right panel: Col1A1 mRNA levels by real-time PCR from CR-PDEL or PMUT mouse livers. Data are mean±S.E from three experimental groups, *p=0.011 vs. oil+EV, #p=0.023 (CR-PDEL) or 0.009 (CR-PMUT) vs. CCI4+EV. (D) Interaction between AKAP12 and HSP47 in desmin-positive HSCs of CRISPR model by PLA staining. Data representative of the PLA/fluorescence count is mean±S.E from 4 PDEL and 3 PMUT experiments. AKAP12-HSP47 PLA: *p=0.018 vs. oil+EV, ⅛p=0.03 vs. CCL4+EV; Desmin: *p=0.02 vs. oil+EV. (E) Top panel: Picosirius red staining of CRISPR-edited livers for collagen as in methods. Three out of 6 PDEL experiments and 3 PMUT experiments are shown. Bottom panel: Hydroxyproline quantification of collagen from6 PDEL and 3 PMUT experiments. *p=2.7X10-5 vs. oil+EV, #p=0.015 vs. CCL4+EV, Sp=0.002 vs. oil+EV or oil+CR.

### HSC-specific phospho-editing of AKAP12 regulates the ER stress response

To determine how HSC-specific AKAP12 phospho-editing reduced overall liver injury and modulated collagen mRNA levels upon CCl4 exposure, we performed proteomics analysis of HSCs and livers isolated from oil+EV, oil+CR, CCl4+EV or CCl4+CR groups to compare the molecular changes under these conditions. Proteomics analysis revealed alterations in several proteins in CCl4 HSCs as well as total liver that were regulated by AKAP12 HSC-specific phospho-editing (Supplementary table S5). Ingenuity pathway analysis (IPA) of these proteins identified two top scoring pathways, the ER stress response and UPR, that were significantly dysregulated by CCl4 and were normalized upon AKAP12 phospho-editing (supplementary table S5). The proteomics analysis of HSCs showed an induction in BIP/GRP78, an ER stress sensor (5), in CCL4-treated group (Figure 7A, Supplementary table S5). We confirmed the proteomics by western blotting. Fold over oil+EV average densitometry is shown below each blot in figure 7 and raw densitometries of each experiment are shown in figure 7-figure supplement 1. HSCs isolated from CCl4 livers showed increased BIP expression (Figure 7B). However, even though the proteomics analysis showed inhibition of CCl4-induced BIP levels by AKAP12 phospho-editing (Figure 7A), we could not confirm this effect by western blotting (Figure 7B). Since BIP is a known collagen chaperone (5), we examined its interaction with collagen in our CRISPR model. BIP exhibited increased interaction with collagen in the CCl4+EV HSCs compared to oil+EV HSCs and AKAP12 phospho-editing strongly inhibited the BIP-collagen interaction in HSCs (Figure 7B). IRE1α, a UPR component that binds to HSP47 and becomes phosphorylated during ER stress (5), exhibited increased interaction with HSP47 in CCL4 HSCs that was inhibited by AKAP12 phospho-editing (Figure 7B). The IRE1α-HSP47 interaction was further confirmed in desmin-positive HSCs of the CRISPR model by PLA staining (Figure 7-figure supplement 2). CCl4-mediated IRE1α phospho-activation (S724 phosphorylation) was strongly inhibited by AKAP12 phospho-editing without a change in total IRE1α levels (Figure 7B). Furthermore, two pathways, P38MAPK and SMAD2/3 that are known to be induced in HSCs through IRE1α activation (30) were also suppressed by AKAP12 phospho-editing (Figure 7B). The proteome of CCl4-exposed livers exhibited increased ER stress and UPR signaling components that were modulated by AKAP12 HSC-specific phospho- editing (Figure 7A, supplementary table S5). BIP levels by western blotting were induced in CCl4 livers and inhibited by AKAP12 phospho-editing, confirming the proteomics result (Figure 7C). Like the proteomics data, we did not find any change in total IRE1α expression. However, phospho-activated IRE1α was suppressed by AKAP12 phospho-editing in total liver (Figure 7C). Since ER stress induces inflammatory signals in different systems (11), we examined whether the HSCs from our CRISPR mouse model exhibited altered inflammatory signaling upon AKAP12 phospho-modulation. Out of the known HSC cytokines, we found the pro-inflammatory cytokine, IL-17, IL-6 and IL-β to be strongly induced in CCl4-HSCs whereas AKAP12-phospho-edited HSCs suppressed their expression (Figure 7D). On the other hand, an anti-inflammatory cytokine, IL-10 was suppressed in HSCs by CCl4 administration, and its expression was restored by AKAP12 phospho-editing (Figure 7D). To examine whether ER stress modulation within activated HSCs was transmitted to other liver cell types, we evaluated crosstalk between HSCs and hepatocytes in a co-culture system where AKAP12 was CRISPR-edited. Co-culture with activated HSCs induced the ER stress response markers BIP and induced IRE1α phosphorylation in hepatocytes compared to co-culture with quiescent HSCs (Figure 7E). Co-culture with activated HSCs in which AKAP12 was phospho-edited (CR) reduced the ER stress signal in hepatocytes compared to activated HSCs alone whereas hepatocytes co-cultured with quiescent HSCs with CR did not exhibit any alteration in ER stress markers compared to WT (Figure 7E). Original blots for figure 7 are shown in Figure 7-source data 1 to 4.

**Figure 7.**
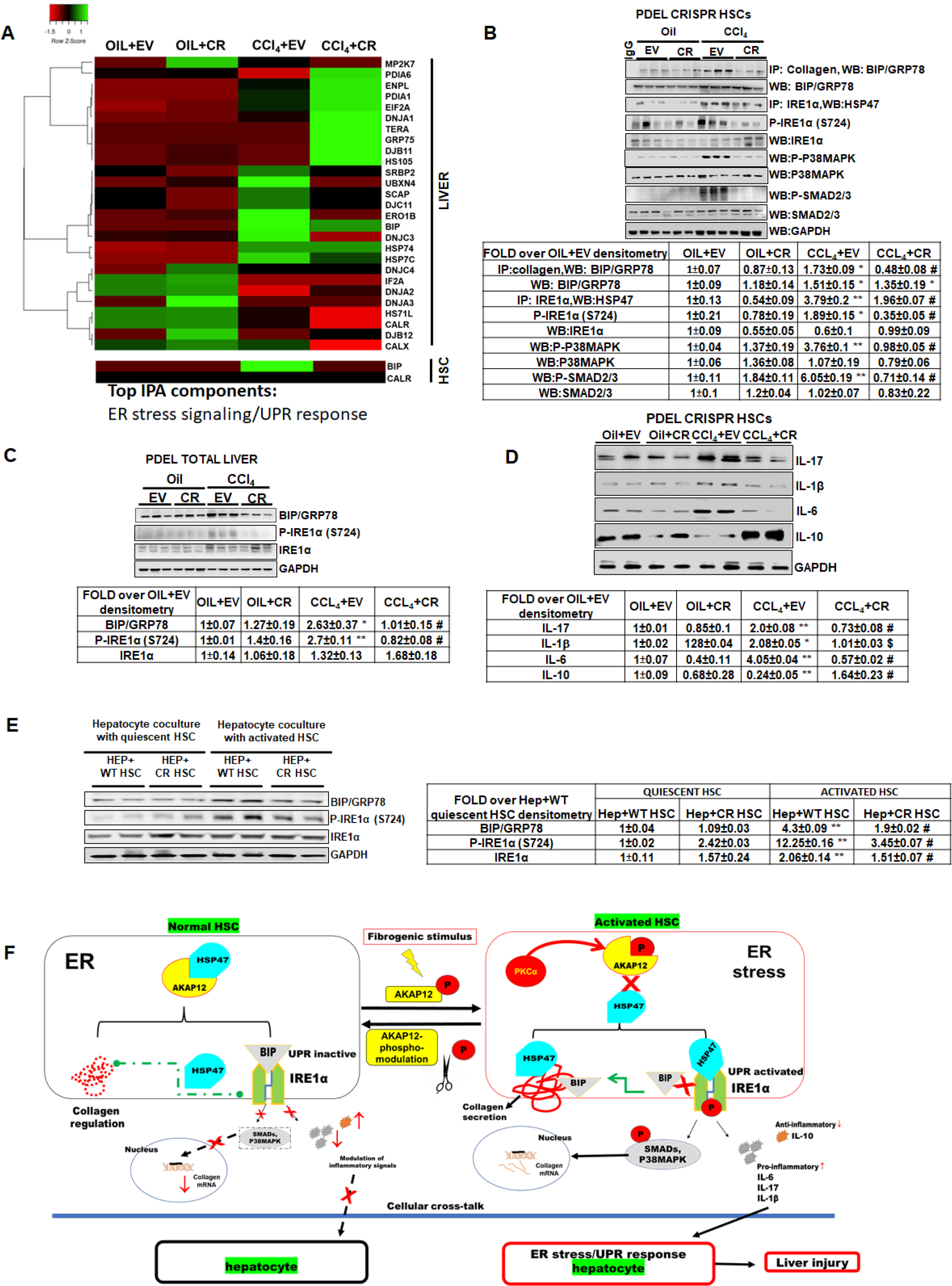
HSC-specífic phospho-editing of AKAP12 regulates the ER stress response. (A) Heat map of total liver and HSCs ER stress√UPR signaling components in four groups, oil+EV, Oil+CR. CCl_4_+EV and CCL_4_+CR. Proteomics data utilized to prepare the heatmap is presented in supplementary table S5>. Alterations in ER stress responsive elements in HSCs from AKAP12 PDEL CRISPR model (B) and total liver (C). GAPDH normalized densitometry represented as fold over oil+EV is meaπ±S.E from 3· experiments. *p<0.05 vs. oil+EV, **p<Q.01 vs. oil+EV, ⅛p<0.01 vs. CCL_a+EV_. Raw densitometry of each experiment is presented in figure S6. (D) Inflammatory cytokine expression in HSCs from oil+EV, Oil+CR, CC∣_i_+EV and CCL_4_+CR groups. GAPDH normalized densitometry represented as fold over oil+EV is mean+S.E from 3 experiments, *p<0.05 vs. oil+EV, **p<θ.θi vs. oil+EV, #p<0.01 vs. CCL_4_+EV, 5p<0.05 vs. CCt_4_+EV. Raw densitometry of each experiment is presented in supplementary figure 6. (E> ER stress response in hepatocytes co-cultured with quiescent or activated WT HSCs with or without AKAP12 PDEL CRISPR editing (CR). GAPDH normalized densitometry represented as fold over hepatocytes+WT quiescent HSCs is mean+S.E from 3 experiments, **p<0.01 vs. h epato cytes+WT quiescent HSC, tfp<0.01 vs. h epato cytes+WT activated HSCs. Raw densitometry of each experiment ispresented in supplementary figure S6. (F) Summary offindings and proposed model of action of phosphσAKAP12. AKAP12 interacts with HSP47 in the ER of normal HSCs and negatively regulates HSP47⅛ collagen-chaperoning activity and its ability to promote ER stress-directed IRE1α branch of UPR signaling. Pro-fit>rogeπicstimuli cause AKAP12”s PKCo-dependent site-specific phosphorylation inhibiting its HSP47 scaffolding activity leading to increased HSP47-collagen chaperoning and UPR signaling via IRE1 α. Consequently, the ER foldase and IRE1α regulator. BIP exhibit increased interaction with collagen. Increased IRE1 α signaling to phospho-5UADs and P38f.1APK induces collagen transcription. Since AKAP12 phospho-edtiπg suppresses HSP47-IRE1α scaffold and IRE1α activation, it also dramatically reduces SMAD and P38MAPK activation and thereby lowers collagen mRNA levels. ER stress∕UPR-liπked inflammatory signals induced upon pro-fibrogenic stimulation are suppressed byAKAP12 phospho-editing in HSCs and total liver. AKAP12 editing also suppresses ER stress response in hepatocyte^HSC cocultures. Blocking AKAP12 phospho-modification inhibits HSC activation, collagen production, fibrosis as well as overall liver injury. We proposea model where ER stressΛJ PR-linked cytokines released fromHSCs may promote ER stress in hepatocytes and ph ospho-edited AKAP12 might inhibit this crosstalk du ring fibrogenesis.

## Discussion

In fibrotic mouse and human livers, HSCs exhibit increased AKAP12 phosphorylation and decreased AKAP12 scaffolding activity towards the collagen chaperone, HSP47. By mapping the phosphorylation events that are altered upon activation of human or mouse HSCs, we have demonstrated that phosphorylation of specific S or T residues of AKAP12 is triggered during HSC activation. Hence, we named these sites as activation-responsive phospho-sites. Out of the 5 activation-responsive phospho-sites, 4 serine residues were confirmed as PKCα substrates. Mutagenesis analysis on recombinant AKAP12 showed that the S687 and S688 were stronger PKCα substrates compared to S676 and S678 because their mutations drastically suppressed phosphorylation. We further observed that phosphorylation of AKAP12 by PKCα suppressed direct binding between AKAP12 and HSP47. Confirming this recombinant data, we observed that silencing *PKCα* in HSCs induced the binding between AKAP12 and HSP47. AKAP12’s known scaffolding activities towards CCND1, PLK1 and PKCα that were identified previously are regulated by its phosphorylation (13, 16 21). We therefore evaluated the role of site-specific phosphorylation in modulating AKAP12’s scaffolding functions in HSCs.

Using a CRISPR-based gene editing approach, we deleted AKAP12’s phosphorylation sites in culture-activated human or mouse HSCs and observed an enhancement in AKAP12’s interaction with HSP47, a strong inhibition of HSC activation (judged by α-SMA levels) and restoration of the quiescent marker, vitamin A that is suppressed in activated HSCs (25). HSP47 resides in the ER (4) and since AKAP12 interacted with HSP47, we evaluated whether it co-localized with HSP47 in the ER and whether the AKAP12-HSP47 scaffold in the ER was affected by CRISPR- editing its phosphorylation sites. The AKAP12-HSP47 scaffold was induced in the ER upon AKAP12 phospho- editing. HSP47’s chaperoning activity towards collagen is highly induced during HSC activation and this allows increased maturation and secretion of collagen (4). Since AKAP12 binds to HSP47, we examined whether this interaction regulated HSP47’s collagen chaperoning function. The ER of activated HSCs stained strongly for the collagen-HSP47 scaffold but AKAP12 phospho-site editing diminished the collagen scaffolding activity of HSP47. Our findings suggest that lack of AKAP12 activation-responsive phosphorylation quenches HSP47’s collagen chaperoning activity and prevents HSC activation.

HSC activation is a hallmark of liver fibrosis. The fact that enhanced phosphorylation of AKAP12 at its activation-responsive phospho-sites promotes HSC activation fueled our hypothesis that site-specific AKAP12 phosphorylation may be involved in promoting liver fibrosis in animal models. To address this hypothesis, we designed CRISPR-AAV vectors to perform gene editing of AKAP12’s activation-responsive phospho-sites specifically in HSCs of mouse liver. This was achieved by expressing the CRISPR causing enzyme, SaCas9 under control of the HSC-specific promoters, GFAP or LRAT (26, 27). Both GFAP and LRAT specifically expressed SaCas9 in HSCs, but GFAP-driven SaCas9 was increased in activated HSCs compared to normal HSCs whereas the reverse was observed with LRAT-Cas9. GFAP promoter activity is induced during HSC activation (31) whereas LRAT expression is known to be suppressed (32). This might have been responsible for the different effects of these two promoters. The AAV particles of serotype 6 were used because AAV6 efficiently transduces activated HSCs in the CCl4 mouse model (33). HSC-specific gene editing of AKAP12 was performed by deleting the DNA region corresponding to the 5 phospho-sites (PDEL). AKAP12 phospho-site editing by this PDEL mechanism strongly inhibited HSC activation, enhanced the AKAP12-HSP47 scaffold, and suppressed the collagen- chaperoning activity of HSP47 leading to decreased collagen production in the liver. To confirm the involvement of AKAP12 phosphorylation at these residues in promoting pro-fibrogenic phenotype, we inhibited phosphorylation at these sites by CRISPR-mediated editing of the S/T residues to A (PMUT). The overall editing efficiency caused by PMUT was lower than that of PDEL in activated HSCs from CCl4-exposed livers. Despite the less efficiency of PMUT editing, it was effective in suppressing the fibrogenic response in the liver, supporting an important role of AKAP12 phosphorylation in regulating the outcome of liver fibrosis. As opposed to fibrotic livers, CRISPR editing in HSCs of normal liver did not alter the molecular identity of the liver. Since normal HSCs do not exhibit phosphorylation of AKAP12 at the activation-responsive phospho-sites, they appear to be unaffected by modulating these sites. This control data reiterates the fact that increased AKAP12 phosphorylation caused by HSC activation has pro-fibrogenic effects.

Apart from suppression of fibrotic parameters, we observed that AKAP12 phospho-modulation in HSCs inhibited collagen mRNA levels and globally suppressed liver injury. Since inhibition of collagen transcription and overall liver injury may not be due to AKAP12’s ability to regulate HSP47’s collagen-chaperoning activity, we searched for additional mechanisms of action of phospho-AKAP12. We performed proteomics analysis in HSCs from our CRISPR model and total liver from the same to identify molecular signals altered by AKAP12 phospho-editing. In HSCs, we identified BIP/GRP78, a regulator of the IRE1α branch of UPR signaling and a known collagen chaperone in the ER (5, 34). Interestingly, a recent interactome study identified HSP47 as a binding partner for IRE1α (5). IRE1α is an ER transmembrane kinase that is kept in inactive state by its binding to BIP. HSP47 activates IRE1α oligomerization and phosphorylation by displacing BIP and triggering the UPR response during ER stress (5). The functional effect of the HSP47-IRE1α interaction on UPR signaling and collagen folding during fibrogenic stimulation in HSCs is undescribed so far. But IRE1α activation caused by ER stress inducers in HSCs is known to enhance collagen transcription as well as collagen protein expression through activation of p38MAPK and SMAD pathways (30). HSCs exhibit ER stress and UPR signaling in response to liver injury stimuli (11, 35). In fact, ER stress appears to be both a cause and effect of HSC activation (11, 36). Since phospho-edited AKAP12 interacted with HSP47 in the ER of HSCs, we wondered whether HSP47-mediated UPR signaling might be regulated by AKAP12. We found that IRE1α-HSP47 interaction (5) was enhanced in CCl4 HSCs and so were downstream pathways known to be enhanced by IRE1α activation in HSCs (phospho-P38MAPK and SMAD2/3) (30). Interestingly AKAP12 phospho-editing suppressed HSP47’s UPR-activating function by quenching the CCl4- mediated IRE1α-HSP47 interaction in HSCs that further inhibited IRE1α phospho-activation preventing downstream P38MAPK and SMAD signaling in these cells. Another component of the UPR signaling we found from proteomics was BIP. BIP is a collagen chaperone that also inactivates IRE1α under basal conditions (5). During ER stress, HSP47 displaces BIP from IRE1α, activating IRE1α-mediated UPR signaling (5). Whether this HSP47-mediated BIP displacement promotes BIP’s activity as a collagen chaperone during HSC activation or liver fibrosis is unclear so far. We hypothesized that AKAP12 by virtue of its increased phosphorylation and lack of scaffolding towards HSP47 may regulate the BIP-IRE1α-HSP47 axis and promote BIP’s collagen chaperoning function. Indeed, we observed increased interaction of BIP with collagen in HSCs of CCL4 livers that was suppressed by AKAP12 phospho-editing. We could not find any interaction between AKAP12 and BIP in HSCs but speculate that loss of AKAP12-HSP47 scaffolding leading to increased HSP47-IRE1α interaction might have released BIP from the IRE1α sites and favored BIP-collagen scaffolding.

Enhanced protein secretion is associated with ER stress and UPR signaling in activated HSCs and is crucial for processing of inflammatory proteins and ECM components upon pro-fibrogenic stimulation (11). Studies in liver and other systems support the role of ER stress in promoting inflammatory signaling (37, 38). Also, inflammatory proteins have a less well described role in promoting ER stress and UPR signaling (38). ER stress is therefore both a cause and consequence of inflammatory signaling (38). Cytokines such as IL-1β are known to be induced in activated HSCs through ER stress (39). Other cytokines known to be expressed by HSCs, IL-17 and IL-6 (40, 41) are prone to modulation by ER stress (42, 43). The anti-inflammatory and anti-fibrotic cytokine, IL-10 expressed by HSCs (44) was recently shown as a target of ER stress in macrophages (45). IL-6 and IL-1β are mediators of ER stress in the liver (46). In pancreatic beta cells, IL-1β is known to induce ER stress in a nitric oxide-dependent manner (47). The anti-inflammatory effect of IL-10 has been shown to block ER stress in intestinal epithelial cells (48). Since our data on AKAP12 suggests that it regulates ER stress pathways in HSCs, we tested whether known inflammatory signals linked to ER stress were also regulated by AKAP12. We found the pro-inflammatory cytokines, IL-17, Il-1β and Il-6 to be induced in CCl4-HSCs whereas AKAP12-phospho-edited HSCs exhibited a strong suppression of these cytokines. On the other hand, the anti-inflammatory cytokine, IL-10 was suppressed in HSCs by CCl4 administration, and its expression was restored by AKAP12 phospho-editing. Literature suggests that inflammatory molecules and UPR signaling may contribute to increased collagen transcription during liver fibrosis.

Pro-inflammatory IL-6 signaling induces collagen transcription (49) and loss of anti-inflammatory signals such as IL-10 inhibit it (50). The IRE1α-directed UPR also induces collagen transcription through increased p38MAPK and SMAD2/3 signaling (30). Since AKAP12 phospho-editing suppressed IRE1α-directed UPR signaling through its association with HSP47 and regulated ER stress-linked cytokines expressed in HSCs, these factors may have contributed to the overall drop in collagen mRNA levels.

Since ER stress/UPR signaling plays a role in enhancing liver injury and the ER stress inducer, tunicamycin is known to induce ALT/AST levels (51), we examined whether AKAP12 HSC-specific editing regulated the liver ER stress response. We found dysregulation of ER stress and UPR-associated components in total liver of CCl4 mice (BIP and other ER foldases such as protein disulfide isomerases, PDIA1 and PDIA6) that were regulated by HSC-specific AKAP12 phospho-editing. Induction of BIP expression in the liver was normalized by AKAP12 phospho- editing. Although the total IRE1α levels were unchanged by CCl4, IRE1α phospho-activation was inhibited by AKAP12 HSC-specific phospho-editing. These results suggest that controlling the ER stress response/UPR signaling within HSCs during pro-fibrogenic stimulation also modulates the same in the whole liver. The phenomenon of ER stress being communicated from stressed cells to other cells within a tissue has been reviewed in the context of cells that produce large amounts of proteins such as immune cells (37). It has also been published that ER stress invokes liver fibrosis primarily due to ER stress within HSCs due to their activation (36). Since hepatocytes are known to be sensitive to CCl4-mediated ER stress (52), we examined whether crosstalk between activated HSCs and hepatocytes in a co-culture system promoted the ER stress response in hepatocytes and whether AKAP12 regulated this crosstalk. Modulating HSC activation through AKAP12 regulated the ER stress response in hepatocytes in culture. Since we observed regulation of ER stress-linked inflammatory cytokine production from HSCs of AKAP12 CRISPR edited livers, we think that inflammatory cytokines from HSCs might transmit ER stress to the whole liver and AKAP12 provides a mode to control these effects during fibrogenesis. A schematic of our findings and their implications in cellular crosstalk during fibrogenesis are summarized in Figure 7F.

In summary, we have identified AKAP12 as a scaffolding partner of HSP47 in normal HSCs that controls HSP47’s collagen chaperoning activity and its interaction with UPR signals in HSCs. Site-specific phosphorylation of AKAP12 occurs during HSC activation and this modification inhibits its interaction with HSP47. This induces HSP47’s collagen chaperoning activity, collagen production and HSP47’s interaction with UPR signaling proteins upon pro-fibrogenic stimulation. Blocking AKAP12 phospho-modification inhibits HSC activation, collagen production, fibrosis as well as overall liver injury possibly via modulation of the ER stress response and inhibition of ER stress-linked inflammatory signals. Structural studies to identify how AKAP12’s activation-responsive phospho-sites interact with HSP47 will facilitate design of small molecules to block AKAP12 phosphorylation and enhance its HSP47 scaffolding activity. Since AKAP12 phospho-modification is not evident in normal HSCs but is induced upon pro-fibrogenic stimulation, AKAP12 phosphorylation may be utilized as a druggable target in liver fibrosis.

## Materials and Methods

### Primary cell isolation and culture

Primary human HSCs purchased from ScienCell, Incorporation, CA) were cultured on plastic dishes for 6 hours (day 0) or further cultured till activation (day 5 or day 7). Mouse HSCs or hepatocytes were isolated from 3-4 months old C57BL/6 mice according to our previously established protocols (24). Mouse HSCs were culture-activated on plastic dishes like human HSCs.

### Phospho-peptide mapping

AKAP12 was immunoprecipitated from HSCs or hepatocytes using an AKAP12 antibody-conjugated protein A/G column (Thermo Scientific). The AKAP12 beads were submitted to Applied Biomics, CA for phospho-peptide mapping. Tryptic peptides were enriched for phospho-peptides and processed for detection of a phospho-site by mass spectrometry. Phosphorylated residues were confirmed by mass spectrometry peak showing the neutral loss of phosphate that was detected from peak shifts on MS/MS spectrum (Table 1, table supplement 1). The observed mass of a phospho-peptide was reduced by 98 Daltons if a single serine/threonine showed a neutral loss of phosphate.

### CRISPR gene editing in cultured HSCs

CRISPR-Cas9 mediated gene editing at the AKAP12 gene locus (exon 3) to delete the region of its activation-responsive phospho-sites was performed by homology-directed repair (HDR). A 22-bp small guide RNA sequence (sgRNA) upstream of a protospacer adjacent motif (PAM- 5’-GTGGAT-3’) recognized by saCas9 (PAM consensus- NNGRRT where N=any nucleotide, R=A or G) (53), was designed and synthesized using the Edit-R CRISPR system (Horizon Discovery, Colorado) (human guide sequence, Supplementary table 1). The CRISPR design tool was used to determine the sgRNA whose sequence is unique compared to the rest of the genome to avoid off-target effects. A donor RNA to delete the phospho-region was designed and synthesized using the Edit-R HDR donor designer system (Horizon) (human PDEL HDR donor, Supplementary table 1). The sgRNA was stabilized by 2’-O-methyl nucleotides and phosphorothioate linkages in the backbone on both the 5′ and 3′ ends and the HDR donor was stabilized by phosphorothioate linkages on both ends to improve functionality during transfection. Cultured cells were co-transfected with a commercially available plasmid, AAV6-GFAP-saCas9, containing the SaCas9 gene under control of the GFAP promoter (Vector Biolabs, PA), sgRNA and HDR donor RNA using the DharmaFECT Duo Transfection Reagent that allows co-transfection of RNA and DNA (Horizon). Cells with transfection reagent alone or SaCas9 plasmid alone+transfection reagent were used as controls. CRISPR designs for mouse HSCs were performed as above for human with mouse guide sequence #1 and mouse PDEL HDR donor (Supplementary table 1). After 48-72 hours of transfection, genomic DNA from human or mouse HSCs was amplified by multiplex PCR using two primers to amplify the region around the deletion site and a third deletion-specific primer to detect HDR-mediated gene editing.

### Gene silencing in activated HSCs

Activated human HSCs (0.3 million cells per well of 6-well plate) were reverse transfected with a universal negative control (Cat# 4404021), PKCα A (Cat# s11092) or PKCα B (Cat# s11094) silencer®select siRNA (Thermo Scientific, IL) using the lipofectamine RNAiMAX reagent as we described previously (24).

### Carbon-tetrachloride (CCl4) injection in mice

12- week-old C57BL/6 male mice were injected intraperitoneally with CCl4 (HPLC grade, Cat#270652, Sigma, diluted 1:3 in mineral oil) or mineral oil (control) at 1 ul/gram body weight bi-weekly for 5-weeks. All procedures for the care and use of mice were approved by the Institutional Animal Care and Use Committee at Cedars-Sinai Medical Center (CSMC).

### CRISPR gene editing in mice

HDR-based gene editing in control or CCl4 mice was performed according to the scheme in figure 4A and B. Two 22-bp sgRNA sequences upstream of a saCas9 PAM (53), were designed using the Edit-R CRISPR system (Horizon Discovery). Off-target analysis for the two sgRNA was performed using the algorithm from the Benchling [Biology Software-(2022) retrieved from https://benchling.com] (supplementary table S6). The two sgRNA sequences were cloned into a single AAV6 vector under the control of a U6 promoter by the cloning service available from Vector builder Inc., IL. An AAV6 vector containing a non-targeting sgRNA was used as an empty vector control (EV). The sequence corresponding to a PDEL or PMUT donor with 500 bp flanking either side of the target region was cloned into a separate AAV6 vector. The PAM sequence in these donors was mutated to prevent re-cleavage by SaCas9 after HDR. The AAV6-GFAP-SaCas9 vector (Vector Biolabs) was used for HSC-specific gene editing. In addition, another AAV6-LRAT-SaCas9 vector was prepared by cloning the mouse LRAT promoter (Accession ID: NM_023624) upstream of SaCas9 (Vector Builder). AAV6 particles of the sgRNA construct, EV construct, PDEL/PMUT donors and GFAP/LRAT-SaCas9 were purified using Vector builder’s AAV production service. For each viral vector, titer was determined by real-time PCR using primers specific for the AAV inverted terminal repeats (ITR). A titer of 1-2X10^13^ genome copies (GC)/ml was achieved for each AAV. All vectors tested negative for mycoplasma contamination. EV or sgRNA vectors along with PDEL or PMUT donors and GFAP or LRAT SaCas9, were injected into tail vein of mice at 10^11^ GC/vector in a volume of 100 ul PBS. Viral vectors were injected into oil or CCl4 mice during the 2^nd^ and 4^th^ week of oil or CCl4 administration (Figure 4B). The HSC specificity of CRISPR was determined by SaCas9 immunofluorescence as described under the immunostaining section. The efficiency of CRISPR editing in HSCs and hepatocytes of gene-edited livers was evaluated by next generation amplicon sequencing. A 298 bp PCR product was amplified from genomic DNA using primers that recognized regions upstream and downstream of the site of AKAP12 deletion or mutation. Amplicons were purified from gels and submitted to Azenta Life Sciences Inc., CA. for performing next generation amplicon sequencing. Briefly, illumina adaptor sequences (FW: 5’- ACACTCTTTCCCTACACGACGCTCTTCCGATCT-3’,REV: 5’- GACTGGAGTTCAGACGTGTGCTCTTCCGA TCT-3’ were added to the amplicons and sequenced by Azenta illumina platform sequencers. The wild type and mutant or deletion mutant reads were counted from each sample and the efficiency of editing was the percentage of edited reads (PDEL or PMUT) versus the total reads. Frequencies of on-target and off-target base changes were analyzed by comparing the target reads to reference reads corresponding to the WT Akap12 amplicon between the two sgRNA sequences (Figure 4A). Within this region, any mismatches other than PDEL or PMUT were considered as off-targets. The mismatches to the reference were observed mainly outside the target region at a frequency of 5% or less (Figure 4G).

### Human tissue array

The human tissue array (Cat# XLiv086-01) in the form of paraffin-embedded tissues was purchased from the human tissue biorepository, US Biolabs Inc. MD. Arrays were stained by immunostaining as described below.

### Real-time RT-PCR

Total RNA from cells or tissues was reverse transcribed to cDNA using M-MLV reverse transcriptase (Nxgen). CDNA was subjected to quantitative RT-PCR using TaqMan probes for mouse *Akap12* and the housekeeping gene, *Gapdh* (mouse) (Life technologies) (24). The PCR profile was: initial denaturation: 95°C for 3 minutes, 45 cycles: 95°C, 3 seconds; 60°C, 30 seconds. The cycle threshold (Ct value) of the target genes was normalized to that of control gene to obtain the delta Ct (ΔCt). The ΔCt was used to find the relative expression of target genes according to the formula: relative expression= 2^-ΔΔCt^, where ΔΔCt= ΔCt of target genes in experimental condition -ΔCt of target gene under control condition.

### Co-immunoprecipitation and western blotting

Total protein extract was processed for immunoprecipitation by incubating 200 µg of pre-cleared protein with with 2 µg of antibody as we described previously (24). Immunoprecipitated protein was processed for western blotting as previously published (24) and developed with Clean-blot™ IP detection reagent (HRP) (Thermo Scientific, IL). Antibodies used for western blotting are listed in supplementary table S7.

### Vitamin A autofluorescence

UV-excited autofluorescence of human HSCs was captured by fluorescence microscopy using a Keyence BZ-X710 inverted fluorescent microscope (Itasca, IL) as we described previously (24).

### Site-directed mutagenesis

An expression vector (pReceiver-WG16) containing the human *AKAP12* gene under control of the T7 promoter was purchased from Genecopoiea, MD and mutated at AKAP12’s activation-responsive sites (S/T to A mutations) using the QuikChange II® site-directed mutagenesis Kit (Stratagene, CA) as we described previously (54). Mutations were detected by sequencing the clones at the Azenta DNA sequencing facility using an *AKAP12* gene-specific primer (5’-GAGAAGGTGTCACTCCC-3’)

### In vitro kinase assay, phostag™ analysis and binding studies

The T7-AKAP12 vector or its mutants were in vitro translated using the non-radioactive TNT® Coupled Transcription/Translation system containing rabbit reticulocyte lysate (RRL) and a biotin-lysyl tRNA according to the manufacturer’s instructions (Promega, WI) to incorporate biotin label into the translated AKAP12 protein. Biotinylated AKAP12 was purified from the RRL components using a biotin-antibody column. Biotinylated AKAP12 or its mutants (5 ul) were used as a substrate for phospho-PKCα in a 25 µl in vitro kinase reaction using 100 ng of active recombinant PKCα enzyme (Millipore-Sigma, MA), 5 µl of a lipid activator (Millipore-Sigma; 20mM MOPS, pH 7.2, 25mM β-glycerolphosphate, 1mM sodium orthovanadate, 1mM dithiothreitol, 1mM CaCl2), 3 µl of Mg^2+/^ATP cocktail (Millipore-Sigma, 20mM MOPS, pH 7.2, 25mM β-glycerophosphate, 5mM EGTA, 1mM Na3VO4, 1mM dithiothreitol, 75mM MgCl2, and 0.5mM ATP) and 2.5 ul of 20 mM Hepes-NaOH buffer, pH7.6. The reaction was carried out at 30°C for 2 hours. The kinase reaction was run on a zinc phostag™ gel containing 15 uM phostag**™** gel (Fujifilm Wako Chemicals, VA) to separate phosphorylated form of AKAP12 from its unphosphorylated counterparts as we described earlier (54). Membranes were probed with streptavidin-HRP (supplementary table S7) to detect biotinylated AKAP12. Biotin antibody was conjugated to protein A/G plus agarose columns using a coupling buffer according to the crosslink immunoprecipitation kit (Thermo Scientific) followed by binding of recombinant biotinylated AKAP12. The columns were treated with recombinant HSP47 protein the absence or presence of active PKCα enzyme. Bound proteins were eluted from the washed column using elution buffer from the crosslinking immunoprecipitation kit (Thermo Scientific) and run on gels along with biotinylated AKAP12 as input and antibody-bound protein A/G beads as IgG controls. Blots were incubated with HSP47 antibody followed by Clean-blot™ IP detection. Reverse IP was done by following the same protocol using HSP47 antibody columns treated with biotinylated AKAP12 followed by detection with streptavidin-HRP. Recombinant HSP47 input was purchased from Prospec protein specialists, NJ.

### Duolink™ proximity ligation assay and immunostaining procedures

For immunocytochemical procedures, cells were fixed with paraformaldehyde and then permeabilized with triton-X 100 before antibody staining. For immunohistochemical analysis, tissues were de-paraffinized and antigen retrieval was performed using the citrate-based antigen unmasking solution (Vector laboratories, CA). For phospho- detection using proximity ligation assay (PLA), primary AKAP12 or phospho-serine (PSer) antibodies (supplementary table 6) were directly conjugated to PLA minus or plus complementary oligonucleotide arms (PLA minus, Catalog no. DUO92010; PLA plus, Catalog no. DUO92009, Millipore-Sigma) according to our previously published protocol (24). To examine protein-protein interactions in cells or tissues, samples were incubated with the antibodies for the interacting targets at 4°C overnight (AKAP12-HSP47, HSP47-collagen). After washing the unbound antibodies, samples were further incubated overnight with secondary antibodies (rabbit or mouse) that were bound to PLA plus or minus complementary probes (Millipore-Sigma, supplementary table S7). The PLA probes were ligated when the proteins were in proximity due to their interaction giving a fluorescent signal as we previously reported (24). To evaluate the localization of interacting partners, co-immunostaining of the PLA signals was done with HSC (desmin) or subcellular compartment (calreticulin ER) marker antibodies. Marker antibodies were detected by Alexa fluor® green rabbit or mouse secondary antibodies (Abcam, supplementary table 6). Co- localization of SaCas9 with desmin or albumin markers in liver tissue was detected by Alexa fluor® secondary antibodies (supplementary table S7). AKAP12 expression in tissues was detected using the mouse HRP/DAB detection immunohistochemistry kit (cat # ab64264, abcam).

### Histopathological examination

Liver sections fixed with 10% neutral formalin were processed for paraffin embedding, sectioning, hematoxylin and eosin (H&E) and picrosirius red staining (collagen) using the services provided by the liver histology core of the University of Southern California research center for liver diseases (NIH grant P30 DK048522).

### Hydroxyproline measurement

The hydroxyproline content of tissue was measured following the protocol from the hydroxyproline assay kit (Cell Biolabs Inc., CA). Briefly, 10 mg of liver tissue was homogenized, and acid hydrolysis was done with 12N HCl. Hydrolyzed samples were treated with chloramine T to convert the hydroxproline to a pyrrole. Ehrlich’s reagent or 4-(Dimethylamino) benzaldehyde added to the pyrrole reacted with it to produce a chromophore whose absorbance could be read at 540-560 nm. The content of hydroxyproline in the tissue sample was determined by comparison to a hydroxyproline standard from the kit that was processed like the unknown sample.

### ALT/AST measurement

ALT and AST levels from plasma of mice were measured with the ALT and AST colorimetric activity assay kits (Cayman Chemical, MI). ALT activity was measured by monitoring the rate of NADH oxidation in a coupled reaction using lactate dehydrogenase (LDH). The NADH to NAD+ oxidation caused a decrease in A340 nm absorbance. The rate of decrease (ΔA340/min) is directly proportional to the ALT activity. AST activity was measured by the rate of NADH oxidation in the presence of malate dehydrogenase. NADH to NAD+ conversion caused a decrease in A340 nm absorbance. Lactate dehydrogenase was added to the AST reaction to prevent interference from endogenous pyruvate in the plasma. The ΔA340/min for both ALT and AST were converted to units/L by dividing the ΔA340 values by the NADH extinction coefficient and multiplying by the sample dilution factor as per the protocol instructions (Cayman).

### Proteomics analysis

Total protein from liver or HSCs was subjected to mass spectrometry-based proteomics analysis by the services of Poochon proteomics solutions, MD. The Nanospray LC/MS/MS analysis of tryptic peptides for each sample was performed sequentially with a blank run between each two sample runs using a Thermo Scientific Orbitrap Exploris 240 Mass Spectrometer and a Thermo Dionex UltiMate 3000 RSLCnano System. Peptides from trypsin digestion were loaded onto a peptide trap cartridge at a flow rate of 5 μL/min. The trapped peptides were eluted onto a reversed-phase Easy-Spray Column PepMap RSLC, C18, 2 μM, 100A, 75 μm × 250 mm (Thermo Scientific, CA) using a linear gradient of acetonitrile (3-36%) in 0.1% formic acid. The elution duration was 110 min at a flow rate of 0.3 μl/min. Eluted peptides from the Easy-Spray column were ionized and sprayed into the mass spectrometer, using a Nano Easy-Spray Ion Source (ThermoScientific) under the following settings: spray voltage, 1.6 kV, Capillary temperature, 275°C. Other settings were empirically determined. Raw data files were searched against mouse protein sequences database using the Proteome Discoverer 1.4 software (ThermoScientific) based on the SEQUEST algorithm. Carbamidomethylation (+57.021 Da) of cysteines was set as fixed modification, and Oxidation / +15.995 Da (M), and Deamidated / +0.984 Da (N, Q) were set as dynamic modifications. The minimum peptide length was specified to be five amino acids. The precursor mass tolerance was set to 15 ppm, whereas fragment mass tolerance was set to 0.05 Da. The maximum false peptide discovery rate was specified as 0.05. The resulting Proteome Discoverer Report contains all assembled proteins with peptides sequences and peptide spectrum match counts (PSM#). The PSM count is a measure of the abundance of the protein.

### Statistical analysis

Western blotting data was quantified by densitometry of blots using the ImageJ software (NIH). PLA staining and immunofluorescence data was analyzed in a blinded manner by two individuals and quantified using ImageJ according to published protocols (55). Scatter bars showing individual experimental points and their means were plotted using GraphPad Prism 9.3.0, GraphPad software. Biologically independent replicates combined from at least three individual experiments were represented as mean ± standard error (Mean±S.E.). Statistical analysis was performed using two-tailed Student’s t-test for paired comparisons and one-way ANOVA (GraphPad Prism) for comparing differences between multiple groups. Significance was defined as P<0.05. Actual p value for each comparison is listed in figure legends.

## Source data figure legends

Figure 1-Source data 1:Raw blots for figure 1A.

Figure 1-Source data 2:Individual images for figure 1B

Figure 1-Source data 3:Raw blots for figure 1C

Figure 2-Source data 1:Raw blots for figure 2B

Figure 2-Source data 2:Individual images for figure 2D.

Figure 3-Source data 1:Raw blots for figure 3A

Figure 3-Source data 2:Raw blots for figure 3B

Figure 3-Source data 3:Raw blots for figure 3C

Figure 4-Source data 1:Original gel for figure 4C and 4E

Figure 5-Source data 1:Six Individual images for PDEL experiment in figure 5B

Figure 6-Source data 1:Raw blots for figure 6A

Figure 6-Source data 2:Raw blots for figure 6B

Figure 6-Source data 3:Raw blots for figure 6C

Figure 6-Source data 4:Six Individual images for PDEL experiment in figure 6E

Figure 7-Source data 1:Raw blots for figure 7B

Figure 7-Source data 2:Raw blots for figure 7C

Figure 7-Source data 3:Raw blots for figure 7D

Figure 7-Source data 4:Raw blots for figure 7E

Figure 2-figure supplement 1-Source data 1:Original gel image

Figure 2-figure supplement 1-Source data 2:Raw blots

Figure 4-figure supplement 1-source data 1:Original gel of supplementary figures 4A and 4C

## Acknowledgements.

This work was supported by NIH grants 1R21ES030534-01A1 (K Ramani) and 1 R21AA027352-01A1 (ML Tomasi, K Ramani).

## Author Contributions

Komal Ramani: Conceptualization, Methodology, Resources, Investigation, Validation, Writing-original draft, funding acquisition; Nirmala Mavila: Methodology, Formal analysis, Writing-review and editing; Aushinie Abeynayake: Methodology, Investigation, Validation; Maria Lauda Tomasi: Writing-review and editing, Formal Analysis; Jiaohong Wang: Investigation, Methodology; Mitchitaka Matsuda: Investigation, Methodology; Ekihiro Seki: Resources, Formal analysis.

---

**Figure 1-figure supplement 1.**
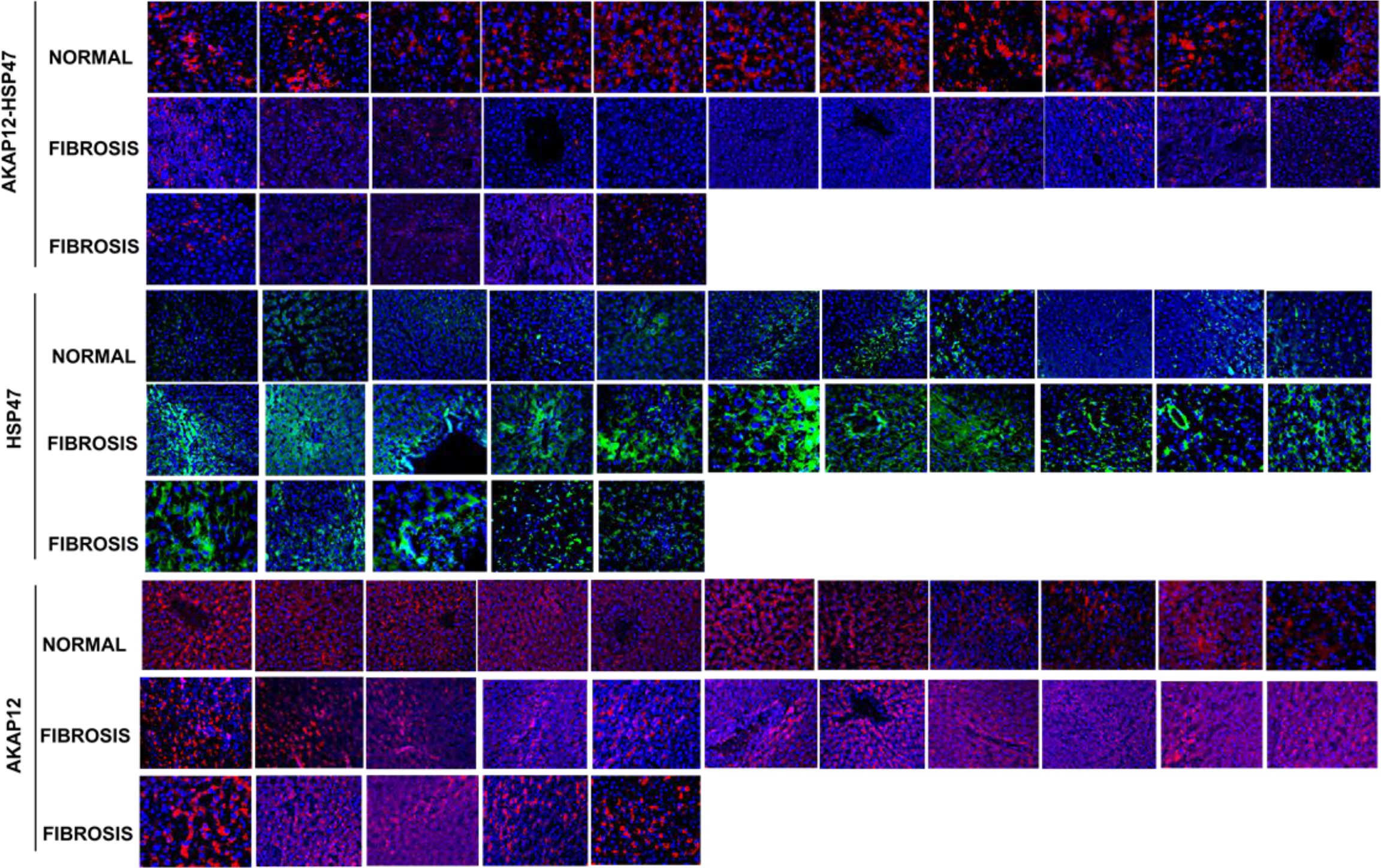
Human liver fibrosis tissue array. The human tissue arrays containing 11 normal livers and 16 liver fibrosis tissues was stained with PLA probes to detect AKAP12-HSP47 interaction and Alexa fluor® probes to detect HSP47 (green) or AKAP12 (red) as described under methods. A representative staining from this complete panel and quantitation by Image J is shown in figure 1D.

**Figure 2-figure supplement 1.**
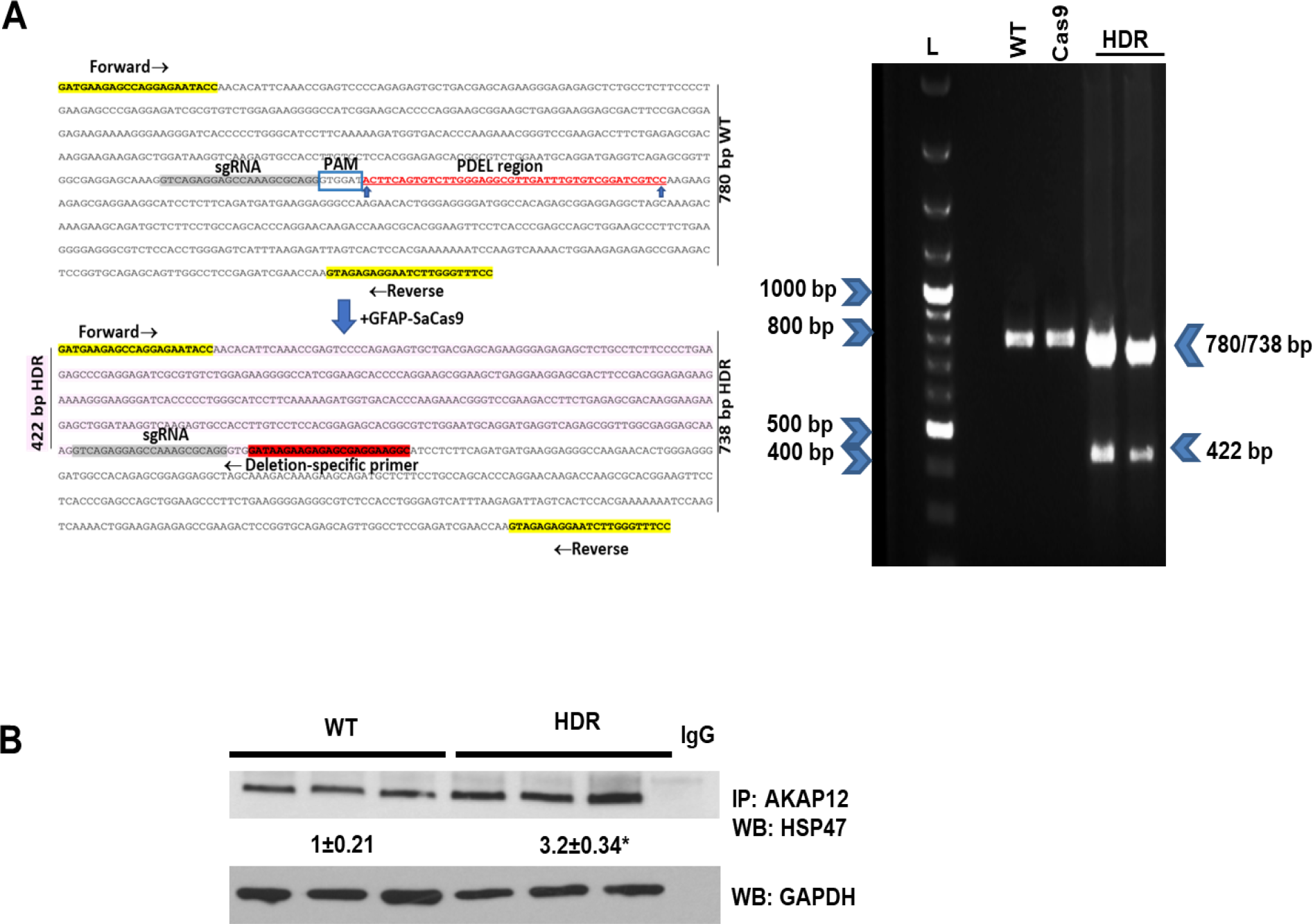
CRISPR-directed editing of AKAP12’s activation-responsive phospho-sites enhances AKAP12’s HSP47 scaffolding activity in mouse HSCs. Activated mouse HSCs were transfected with CRISPR reagents and GFAP-Cas9 vector to cause CRISPR-directed HDR as described under methods. Untransfected (WT) or cells with Cas9 alone were used as controls. (A) CRISPR editing at the AKAP12 locus (left panel) was confirmed by performing PCR (right panel) using primers that specifically detected the edited region as listed in supplemental table S1. A representative image from 3 experiments is shown. (B) CRISPR-edited (HDR) or WT cells were assessed for AKAP12-HSP47 interaction by co-immunoprecipitation-immunoblotting. Data represented as GAPDH normalized densitometry is mean±S.E from 3 experiments. *p=0.03 vs. WT.

**Figure 4-figure supplement 1.**
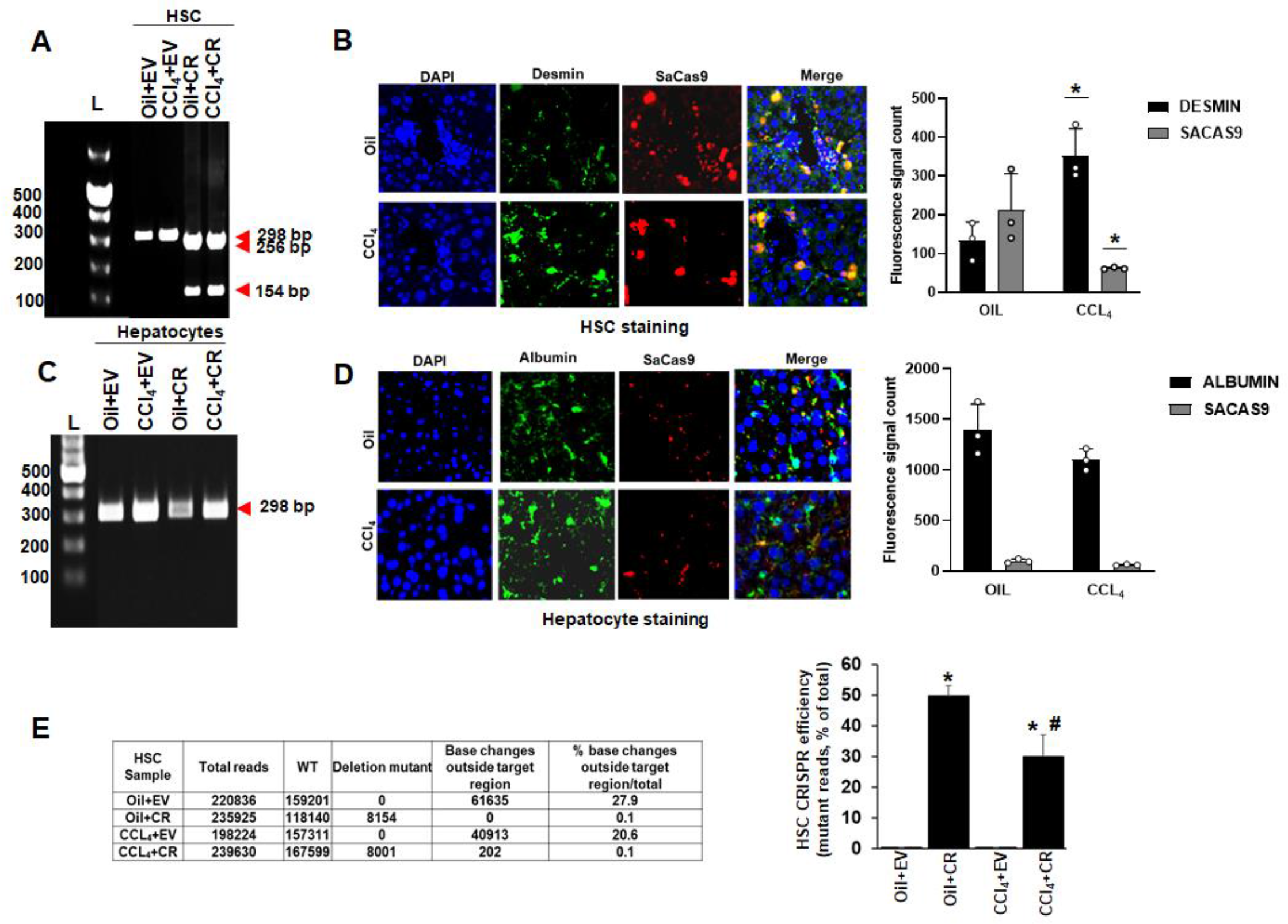
In vivo gene editing of the *Akap12* region corresponding to its activation-responsive phospho- sites in the CCl4 mouse model using LRAT-Cas9. (A) Specificity of PDEL CRISPR for HSCs using LRAT-SaCas9 was evaluated by multiplex PCR amplification of genomic DNA from HSCs using a PDEL-specific primer and two primers around the PDEL primer site (Supplementary table 1). A representative gel image from 3 experimental groups is shown. (B) Specificity of HSC CRISPR was ascertained by co-localization of SaCas9 with desmin-positive HSCs in oil or CCl4-treated groups. Data is representative of three experiments (400X magnification). Desmin staining: *p=0.011 vs. oil; SaCas9 staining: *p=0.049 vs. oil. (C) Multiplex PCR of hepatocytes genomic DNA as in ‘A’ above did not show PDEL specific amplicons. A representative gel image from 3 experiments is shown. (D) SaCas9 co-localization with albumin-positive hepatocytes was insignificant in oil or CCl4 livers compared to HSCs in ‘B’ above. Data are representative of 3 experiments (400X magnification). (E) The efficiency of CRISR in HSCs was evaluated by next generation amplicon sequencing using a 298 bp PCR amplicon derived from genomic DNA. Total amplicon reads, WT reads, and PDEL reads within the target region or base changes outside the target region from each experimental group are shown. The CRISPR editing efficiency represented by the percentage of mutant reads versus total is mean±S.E from 3 PDEL experiments. *Oil+CR or CCL_4_+CR-PDEL p=0.002 vs. Oil+EV, #CCl_4_+CR-PDEL p=0.001 vs. CCl4+EV.

**Figure 5-figure supplement 1.**
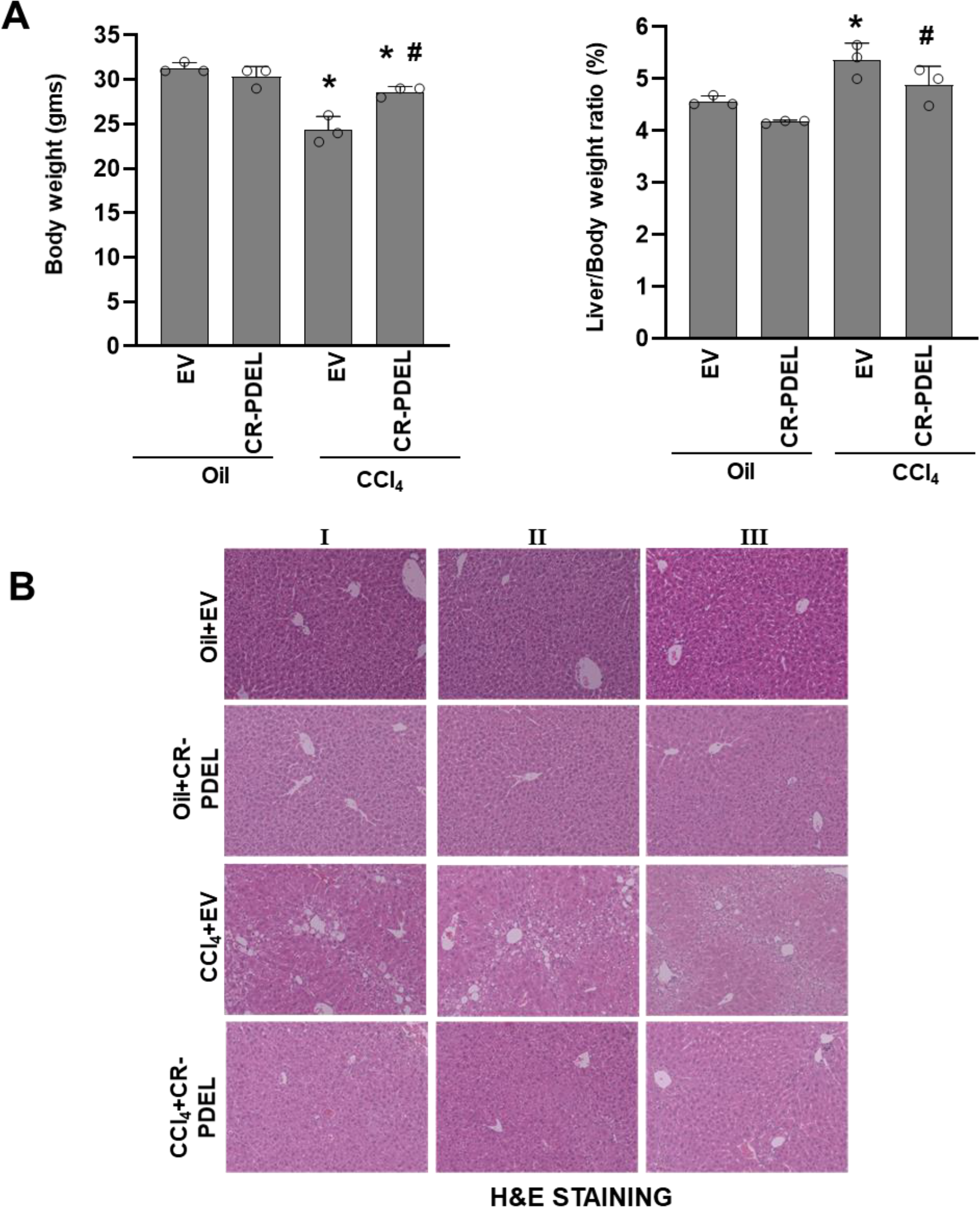
Phospho-editing of AKAP12 using LRAT-Cas9 regulates liver fibrosis in the CCl4 mouse model. (A) Gross changes in mouse body weight (left panel) and liver/body weight ratio (right panel) after PDEL LRAT-Cas9-mediated CRISPR editing of AKAP12’s phospho-sites under oil or CCl4 treatment conditions. Mean±S.E from 3 PDEL experiments. CCl4+EV: *p=0.0018 vs. Oil+EV; CCL4+CR-PDEL: *p=0.0048 vs. oil+EV or #p=0.01 vs. CCl4+EV. **(B)** Histological evaluation of CRISPR-edited livers by H&E staining as described under methods. Three PDEL experiments with LRAT-Cas9 are shown.

**Figure 7-figure supplement 1.**
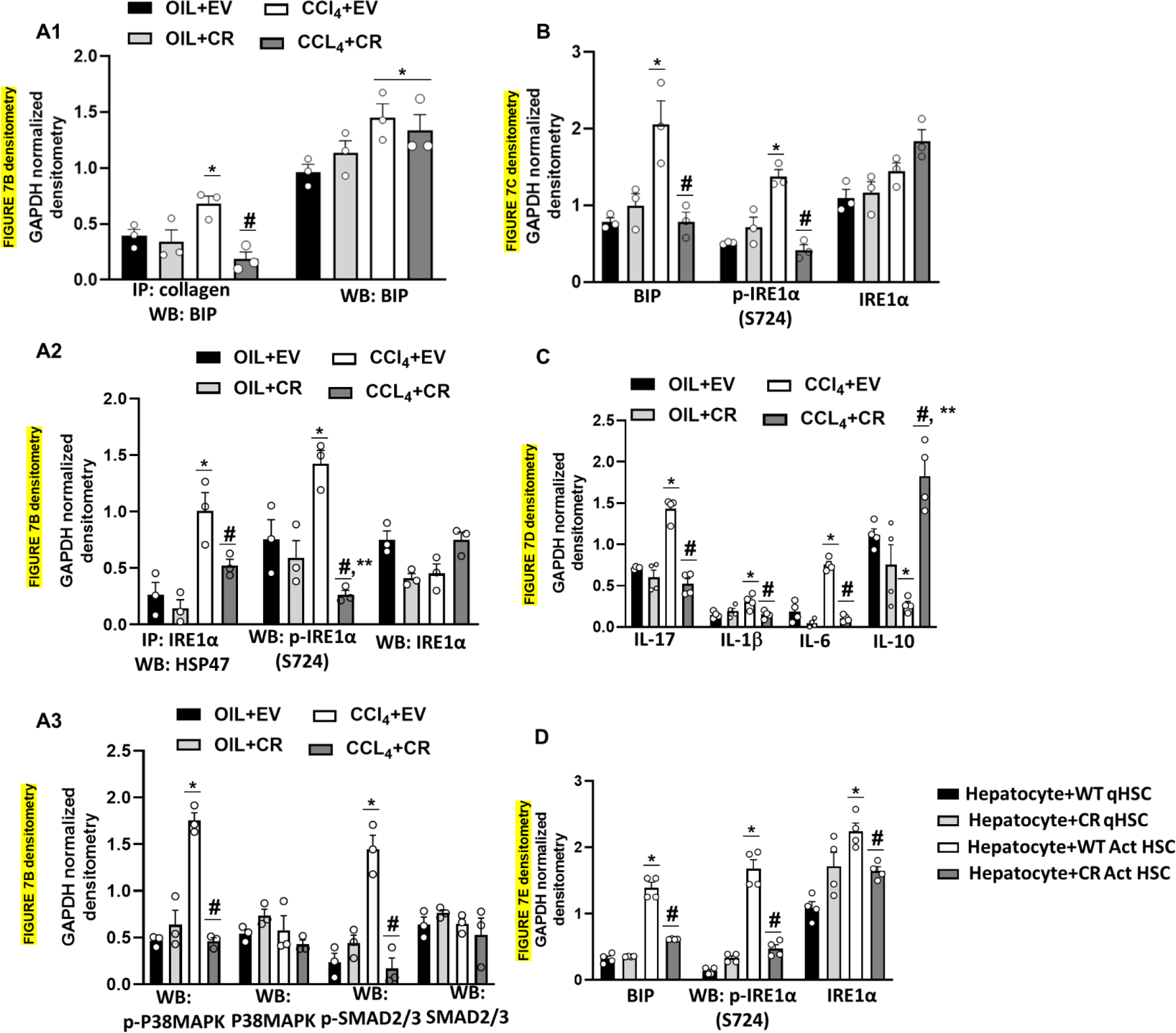
Densitometric quantification of blots from. Figure 7B **to 7E**. (**A1**). PDEL-CR-HSC. IP: collagen, WB: BIP: *p=0.036 vs. Oil+EV, #p=0.006 vs. CCl4+EV; WB: BIP: *p=0.05 vs. Oil+EV. (**A2**) PDEL-CR-HSC. IP: IRE1α, WB: HSP47: *p=0.019 vs. Oil+EV, #p=0.048 vs. CCl4+EV; P- IRE1α (S724): *p=0.03 vs. Oil+EV, #p=0.0007 vs. CCl4+EV, **p=0.049 vs. Oil+EV. (**A3**) PDEL-CR-HSC. p-P38MAPK: *p=0.00015 vs. Oil+EV, #p=0.000157 vs. CCl4+EV; p-SMAD2/3: *p=0.0026 vs. Oil+EV, #p=0.0026 vs. CCl4=EV. **(B**) CR-PDEL Total liver. BIP: *p=0.014 vs. Oil+EV, #p=0.018 vs. CCl4+EV; P- IRE1α (S724): *p=0.0006 vs. Oil+EV, #p=0.0009 vs. CCl4+EV. (**C**) PDEL-CR-HSC. IL17: *p=5.4X10-5 vs. Oil+EV, #p=8.4X10- 5 vs. CCl4+EV; il-1β: *p=0.018, #p=0.02 vs. CCl4+EV; IL-6: *p=0.0002 vs. Oil+EV, #p= 5.0X10-6 vs. CCl4+EV; IL-10: *p=7.14X10-5 vs. Oil+EV, #p=0.0003 vs. CCl4 +EV, **p=0.016 vs. Oil+EV. (**D**) BIP: *p=2.2X10-5 vs. hepatocyte+WT qHSC, #p=7.8X10-5 vs. hepatocyte+WT Act HSC; P- IRE1α (S724): *p=3X10-5, vs. hepatocyte+WT qHSC, #p=0.0002 vs. hepatocyte+WT Act HSC; IRE1α: *p=0.0003 vs. hepatocyte+WT qHSC, #p= 0.005 vs. hepatocyte+WT Act HSC.

**Figure 7-figure supplement 2.**
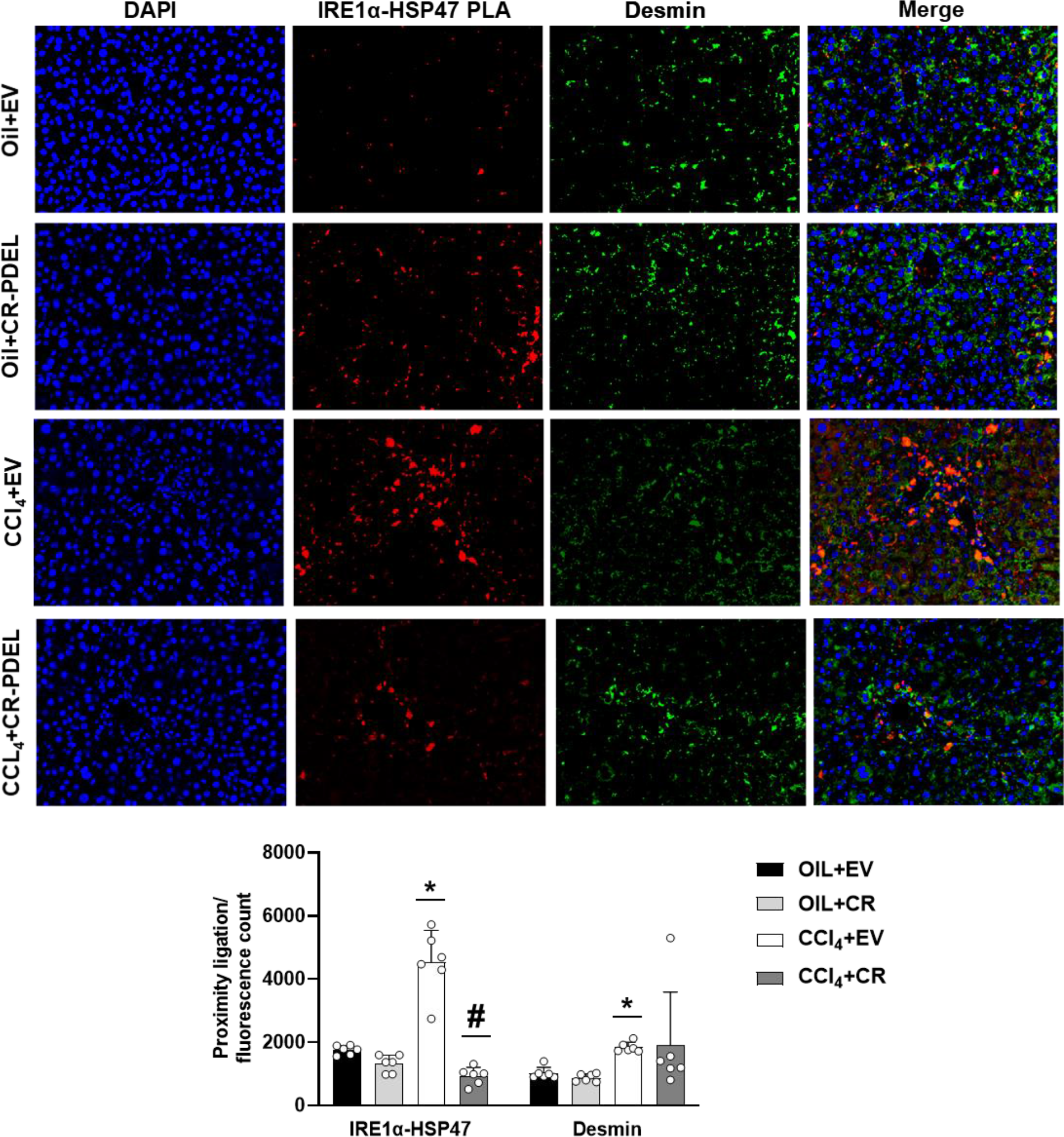
**Interaction between IRE1α and HSP47 in desmin-positive HSCs of CRISPR-PDEL model by PLA staining**. Data representative of the PLA/fluorescence count is mean±S.E from 6 experiments (200X magnification). IRE1α-HSP47 PLA: *p=0.00006 vs. Oil+EV, #p=0.00003 vs. CCl4+EV; Desmin: *p=9.57X10^-6^ vs. oil+EV.

**Supplementary Table S1.**
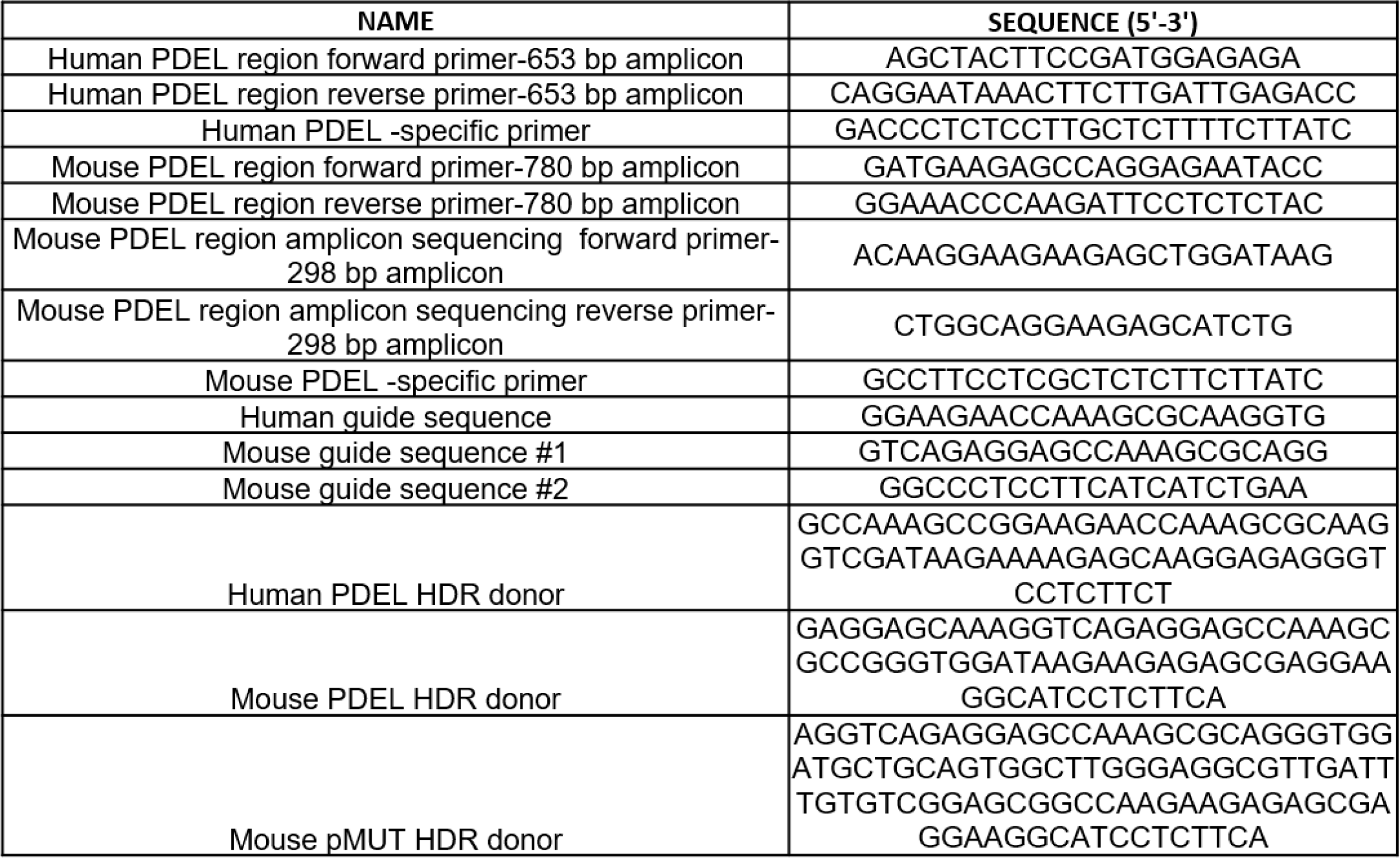
Sequences of PCR primers, CRISPR guides and CRISPR donors

**Supplementary Table S2.**
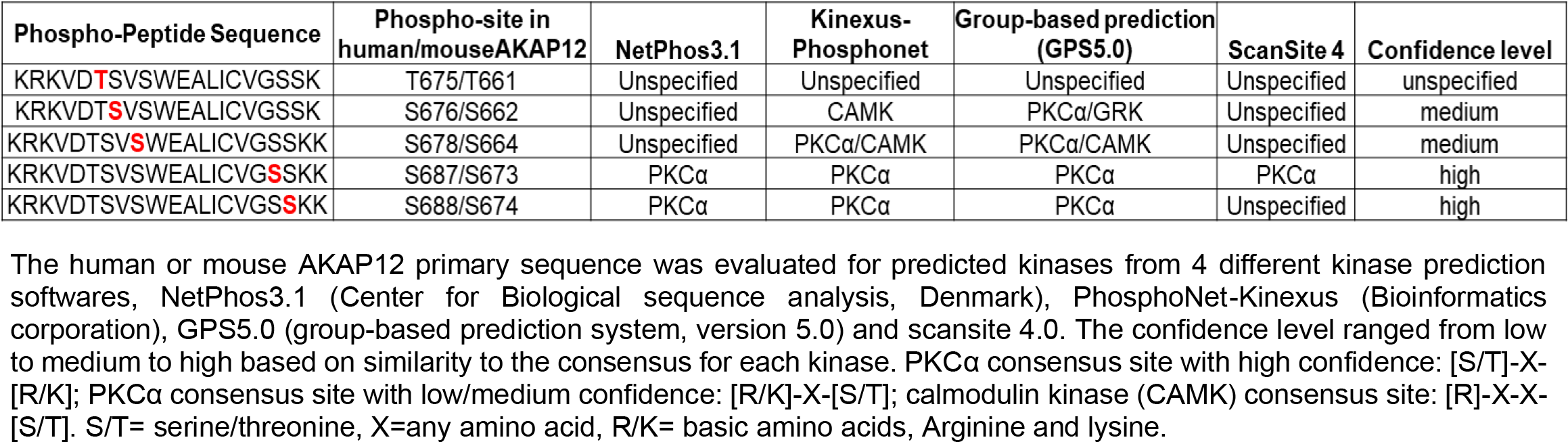
Kinase-prediction for AKAP12’s activation-responsive phospho-sites

**Supplementary Table S3.** Datasets of next generation amplicon sequencing (NGS) from PDEL CRISPR mouse model. Genomic DNA of HSCs isolated from oil or CCl4 injected mice treated with AKAP12 PDEL CRISPR+GFAP-Cas9 or LRAT-Cas9 were submitted for NGS to Azenta Life Sciences as described under methods. Hepatocytes from PDEL CRISPR+GFAP-Cas9 were also processed as above for NGS. Representative raw reads of WT, deletion or base changes are shown for each data set and summarized in the first summary tab of the excel.

**Supplementary Table S4. Datasets of next generation amplicon sequencing (NGS) from PMUT CRISPR mouse model.** Genomic DNA of HSCs or hepatocytes isolated from oil or CCl4 injected mice treated with AKAP12 PMUT CRISPR+GFAP-Cas9 were submitted for NGS to Azenta Life Sciences as described under methods. Representative raw reads of WT or base changes are shown for each data set and summarized in the first summary tab of the excel.

**Supplementary Table S5. Proteomics analysis of total liver and HSCs from CRISPR PDEL mouse model.** Total protein from the liver or HSCs of AKAP12 PDEL CRISPR+GFAP-Cas9 mice was subjected to mass spectrometrybased proteomics analysis as described under methods. Proteomics dataset of whole liver, ER stress/UPR components of the liver and HSCs is shown. The summary tab in the excel explains each dataset.

**Supplementary Table S6.**
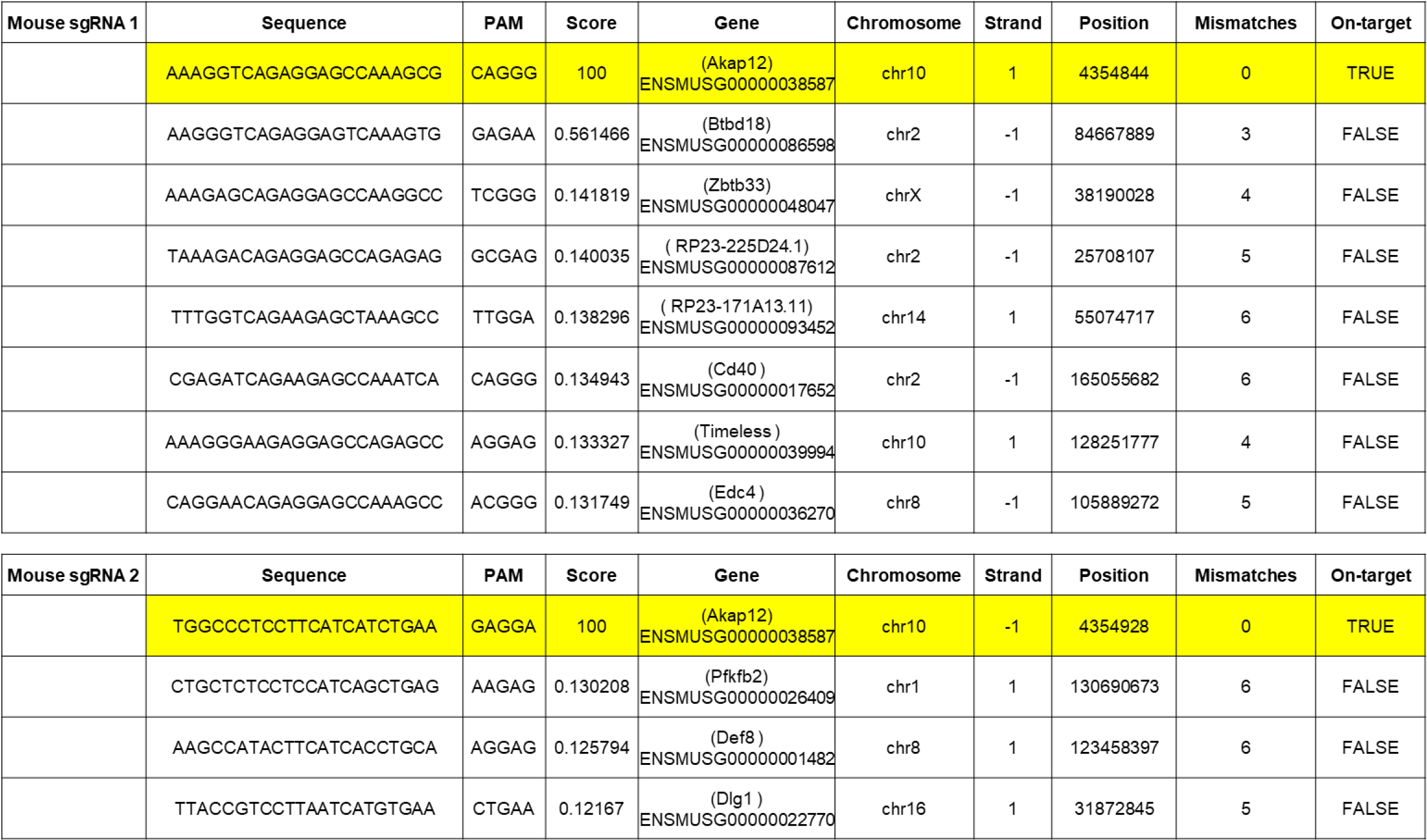
Mouse sgRNA off-target analysis

**Supplementary Table S7.** Antibodies used in this study

| NAME | SUPPLIER | CATALOG# | CLONE | APPLICATION |
| --- | --- | --- | --- | --- |
| Anti-AKAP12 antibody | Abcam | ab49849 | JP74 monoclonal | Primary antibody: co-immunoprecipitation, western blotting, immunostaining, PLA |
| Anti-Phosphoserine antibody | Abcam | ab9332 | Polyclonal IgG | Primary antibody: PLA |
| Anti-α-SMA antibody | Abcam | ab5694 | Polyclonal IgG | Primary antibody: western blotting |
| Anti-PKCα antibody | Genetex | GTX130453 | Polyclonal IgG | Primary antibody: western blotting |
| Anti-Collagen I alpha (1) antibody | Novus Biologicals | NB600-450 | COL-1 monoclonal | Primary antibody: western blotting, PLA |
| Anti-HSP47 antibody | Novus Biologicals | MAB9166-100 | 950806 monoclonal | Primary antibody: co-immunoprecipitation, western blotting, immunostaining, PLA |
| Anti-Biotin antibody | Abcam | ab53494 | Polyclonal IgG | Primary antibody: western blotting |
| Anti-GAPDH antibody | Proteintech | 10494-1-AP | Polyclonal IgG | Primary antibody: immunoprecipitation, western blotting |
| Anti-SaCas9 antibody | Genetex | A01951 | 11C12 monoclonal | Primary antibody: Immunostaining |
| Anti-desmin antibody | Proteintech | 16520-1-AP | Polyclonal IgG | Primary antibody: Immunostaining |
| Anti-albumin antibody | Proteintech | 16475-1-AP | Polyclonal IgG | Primary antibody: Immunostaining |
| anti-IRE1α antibody | Proteintech | 27528-1-AP | Polyclonal IgG | Primary antibody: PLA, western blotting |
| anti-Phospho-IRE1α (S724) antibody | Abcam | ab226974 | Polyclonal IgG | Primary antibody: western blotting |
| Anti-phospho-Smad2 (Ser465/467)/Smad3 (Ser423/425) | Cell Signaling | 8828 | Polyclonal IgG | Primary antibody: western blotting |
| Anti-SMAD2 antibody | Proteintech | 12570-1-AP | Polyclonal IgG | Primary antibody: western blotting |
| Anti-SMAD2 antibody | Proteintech | 25494-1-AP | Polyclonal IgG | Primary antibody: western blotting |
| anti-BIP/GRP78 antibody | Proteintech | 11587-1-AP | Polyclonal IgG | Primary antibody: co-immunoprecipitation, western blotting |
| anti-IL1β antibody | Proteintech | 26048-1-AP | Polyclonal IgG | Primary antibody: western blotting |
| anti-IL6 antibody | Proteintech | 21865-1-AP | Polyclonal IgG | Primary antibody: western blotting |
| anti-IL17 antibody | Proteintech | 66148-1-Ig | 1B3D5 monoclonal | Primary antibody: western blotting |
| anti-IL10 antibody | Proteintech | 20850-1-AP | Polyclonal IgG | Primary antibody: western blotting |
| Anti-calreticulin antibody | Abcam | ab92516 | EPR3924 monoclonal | Primary antibody: Immunostaining |
| Clean-Blot™ IP Detection (HRP) | Life technologies | 21230 | Secondary antibody | Detection: co-immunoprecipitation-immunoblot |
| Streptavidin-HRP | Cell Signaling | 3999 | Secondary antibody | Detection: Biotin western blots |
| Goat anti rabbit IgG H&L (Alexa Fluor® 488 green) | Abcam | ab150077 | Secondary antibody | Detection: immunofluorescence |
| Goat Anti-Mouse IgG H&L (Alexa Fluor® 488 green) | Abcam | ab150113 | Secondary antibody | Detection: immunofluorescence |
| Goat Anti-Mouse IgG H&L (Alexa Fluor® 647 far red) | Abcam | ab150115 | Secondary antibody | Detection: immunofluorescence |
| Goat Anti-Rabbit IgG H&L (Alexa Fluor® 647 far red) | Abcam | ab150079 | Secondary antibody | Detection: immunofluorescence |
| Duolink® In Situ PLA® Probe Anti-Mouse PLUS | Millipore-Sigma | DUO92001 | Secondary antibody | Detection:PLA |
| Duolink® In Situ PLA® Probe Anti-Mouse MINUS | Millipore-Sigma | DUO92004 | Secondary antibody | Detection:PLA |
| Duolink® In Situ PLA® Probe Anti-Rabbit PLUS | Millipore-Sigma | DUO92002 | Secondary antibody | Detection:PLA |
| Duolink® In Situ PLA® Probe Anti-Rabbit MINUS | Millipore-Sigma | DUO92005 | Secondary antibody | Detection:PLA |

